# Baobab isotope records and rainfall forcing in southwest Madagascar over the last 700 years

**DOI:** 10.1101/2025.08.15.670475

**Authors:** Estelle Razanatsoa, Lindsey Gillson, Grant Hall, Malika Virah-Sawmy, Stephan Woodborne

## Abstract

Highly resolved climate records for Madagascar are scarce but are essential for understanding of rainfall drivers over time and assessing the risks and likely trajectories of future climate change. We measured variation in the carbon isotopes of baobabs (*Adansonia* spp.) which reflect rainfall in southwest Madagascar. The record indicates a decreasing trend of rainfall over the last 700 years with high variability at a centennial-scale. The duration of wetter periods decreased over time with the wettest periods between 1350 – 1450 CE, after the onset of the Little Ice Age, while the driest period occurred between 1600 – 1750 CE, during the Maunder Minimum. The results suggest that decadal to centennial rainfall variability in southwest Madagascar is dominated by tropical forcing rather than subtropical forcing. Wetter periods are regulated by the movement and migration of easterly winds linked to the Intertropical Convergence Zone, while dry periods are influenced by the effect of the Pacific Decadal Oscillation linked to the El Niño Southern Oscillation and the sea surface temperature variation in the Southwestern Indian Ocean. The Southern Annular Mode is significantly correlated with the record, but its effect was only visible at the beginning of the record around 1300 CE. This evidence provides a new understanding of rainfall across southern Africa and the interaction of global forcing with regional factors. Further investigation is required to improve tree chronology from Southern Hemisphere and understand the migration of the westerlies and its potential future effect on the rainfall in Madagascar. Understanding the interplay between tropical and other rainfall forcings will be essential in assessing likely scenarios of resilience, and adaptive capacity of social-ecological systems in Madagascar.

## 1. INTRODUCTION

The African continent has been tangibly affected by climate change in the last decades, manifested by increasing temperatures and rainfall variability (IPCC, 2021; Baptista et al., 2022; Weathering Risk, 2023; Onyeaka et al., 2024). In parts of southern Africa, decreased rainfall with more pronounced and recurrent severe drought events are expected in the near future (e.g Ingram & Dawson, 2005; Heland & Folke, 2014; Hoscilo et al., 2015; Baudoin et al., 2017; Serele et al., 2020; IPCC, 2021). Understanding rainfall drivers requires knowledge of local and regional synoptic changes that occur at decadal to centennial scales (Scroxton et al., 2017). Madagascar plays a role in the regulation of climate across southern Africa through its topographic influence on the Mozambique Channel Trough (MCT, Barimalala et al., 2018). Madagascar has an east-west rainfall gradient associated with its topography which reduces the influence of easterly trade winds that bring moisture from equatorial Indian Ocean (Jury, 2016; Fig. 1). A north-south rainfall gradient derives from the north-westerly monsoon as the ITCZ crosses Madagascar to 20°S during the austral summer (Donque, 1975; Jury & Huang, 2004; Tadross et al., 2008; Scroxton et al., 2017). While the southwest region derives moisture from the monsoon and tropical cyclone events in the South Indian Ocean (east Madagascar), and the Mozambique Channel (west-southwestern Madagascar) (Ho et al., 2006), it is the driest area on the island with <600 mm of rainfall per year with recurrent drought events (Ganzhorn, 1995; Tadross et al., 2008). The southern region of Madagascar has experienced limited rainfall at least in the last few decades (Serele et al., 2020; Harrington et al., 2021; Harrington et al., 2022; Otto et al., 2022). The region is also vulnerable to future global climate changes as simulations suggest more severe and longer dry seasons in the tropics, and an average global rise in temperature of 1.5°C estimated to be reached by 2030 (Christensen et al., 2007; IPCC, 2021; Barimalala et al., 2021). This forecast is associated with large uncertainty due to a lack of understanding of seasonal, annual, decadal, and centennial rainfall trends, and this prevents proper assessment of adaptation and hazard reduction in the area (Serele et al., 2020).

**Fig. 1:**
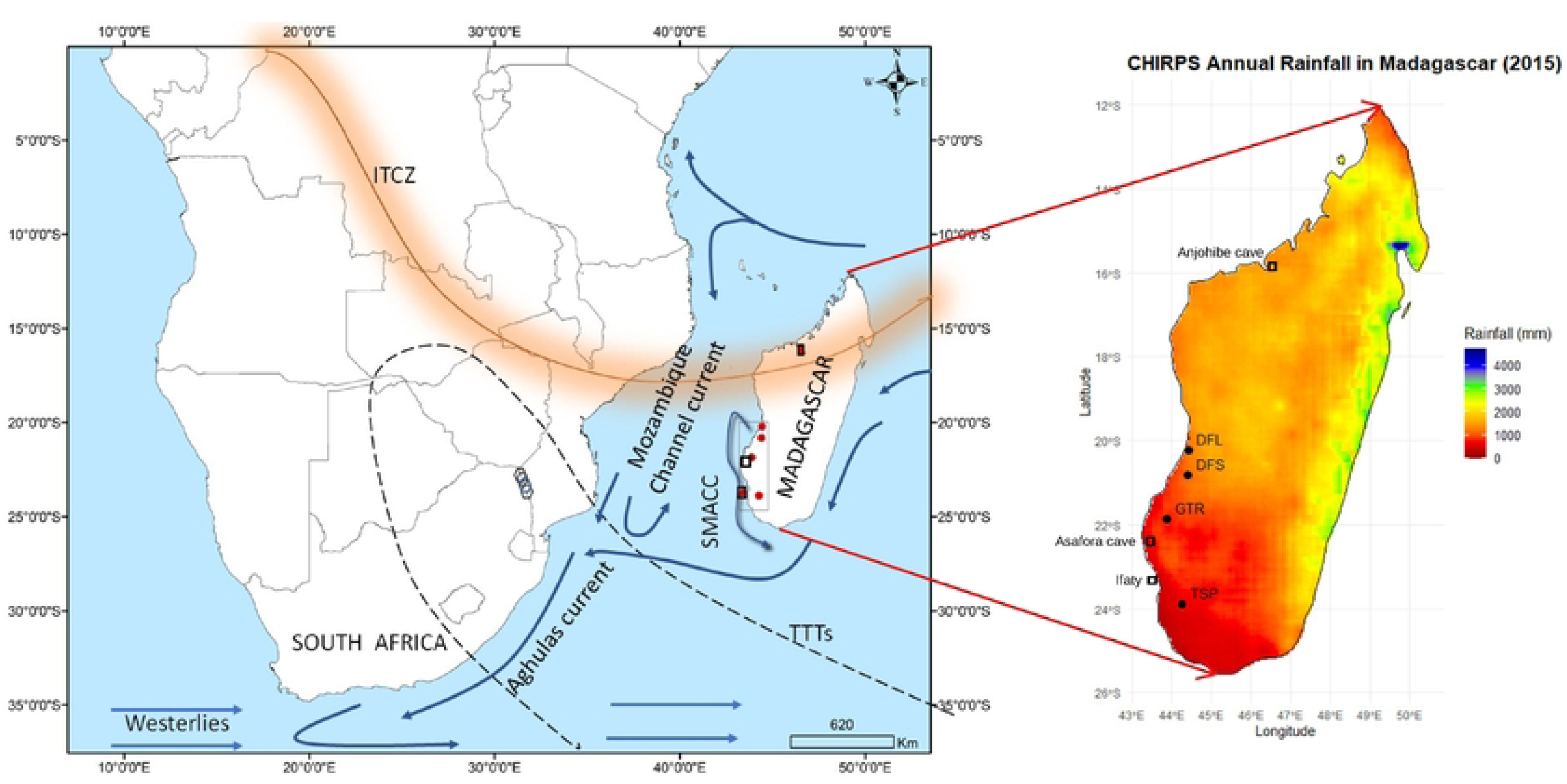
(A) Location of the four baobabs that were analysed from southwest Madagascar (red) and austral summer synoptic features in southern Africa, including the Inter Tropical Convergence Zone (ITCZ) during the austral summer, Tropical Temperate Troughs, Southwest Madagascar Coastal Current (SMACC), Agulhas Current and Mozambique Channel Current. The locations of the SST coral records from Ifaty in southwest Madagascar and the speleothem record from Anjohibe cave in northwest Madagascar, stalagmites records from Asafora cave and additional tree records (hexagons) from southern Africa are shown. (B) Madagascar rainfall gradient with the four trees (circle) and other proxy records used in this study (rectangle).

On an annual to decadal scale, droughts in southern Africa have been associated with El Niño Southern Oscillation (ENSO; Baudoin et al., 2017). Austral summer rainfall is controlled primarily by the seasonal interplay between subtropical high-pressure systems and the migration of easterly flows associated with the Intertropical Convergence Zone (ITCZ) (Chase & Meadows, 2007; Sachs et al., 2009). The position and intensity of tropical-temperate troughs (TTTs) and their associated cloud bands that form in response to this interplay (Mason & Jury, 1997) are closely linked to sea surface temperature (SST) conditions in the Southwest Indian Ocean (SWIO) (Reason & Mulenga, 1999; Nash, 2017, Fauchereau et al., 2009). These synoptic dynamics are derived from observational data, longer term data are needed to understand trends and the interplay between the spatial dynamics of rainfall and the different drivers at decadal and centennial scales. This is especially true for Madagascar where rainfall variability is intrinsically high; there is a high dependence in rain-fed agriculture; and environmental vulnerability is high.

The limited climate analyses from the island show drying trends at millennial, centennial to decadal scales (Ganzhorn, 1995; Tadross et al., 2008; Virah-Sawmy et al., 2016). ENSO and the seasonality of the ITCZ are essentially tropical forcing mechanisms, but recent evidence suggests that under glacial climate forcing, there is an influence from the sub-tropics that is related to the latitudinal position of the southern hemisphere westerlies (Hahn et al., 2021). The South Annular Mode (SAM) is a position index of the westerly vortex, but little is known about its past and current effect on rainfall in the region (Nash, 2017) although it is known to affect circulation on weekly to centennial time scales (e.g., Trenberth, 1979; Thompson & Wallace, 2000).

### Paleoclimate proxy records

Paleoclimate records contain information at various temporal and spatial scales, which enables the interpretation of trends, variability, and the underlying processes within the climate system. Paleoclimate proxy records derived from ice cores (e.g. Svensson et al., 2006; Thompson et al., 2003), lake and wetland sediments (e.g. Neukom & Gergis, 2011; Verschuren, 2003), and speleothems (e.g. Holmgren et al., 1999; Voarintsoa et al., 2017; Scroxton et al., 2017) provide a deep time record of global-scale climate forcing, while coral reefs (e.g. Zinke et al., 2014) and tree rings (e.g. Cook et al., 2002; Helama et al., 2005) yield shorter, more localised records of climatic responses. In temperate environments, abundant records of tree ring widths and varved sediments growth can be used to infer rainfall regimes, but records with similar resolution are rare in tropical ecosystems. Recently the possibility of reconstructing past climate variability using carbon isotope ratios (*δ*^13^C) from the growth rings of subtropical trees has emerged (Robertson et al. 2006; Woodborne et al., 2015; 2016). This increases the potential for paleohydrological reconstruction in the subtropics, and the approach holds the potential to address the climate data deficit in Madagascar which has numerous species of long-lived, dry adapted trees such as the baobab (Baum & Baum, 1995; Baum et al., 1998; Razanamaro et al., 2015).

### Carbon isotope in tree rings as a climate proxy

The use of *δ*^13^C as a climate proxy in tree growth is based on environmental regulation of carbon isotope fractionation during photosynthesis. Fractionation is controlled by stomatal conductance, which manifests in the ratio of leaf internal CO_2_ concentration and the atmospheric CO_2_ concentration (*c_i_*/*c_a_*). Many environmental factors control stomatal conductance (Farquhar et al., 1982; Tieszen, 1991), but in ecosystems where water stress is the main control, reducing the *c_i_*/*c_a_*and increases the *δ*^13^C values of plants (more positive) in dry conditions. In low rainfall regions, water stress responses in trees relate to edaphic water availability which is directly linked to rainfall (Farquhar et al., 1982). In the southern African subtropics, the *δ*^13^C value of a specific growth ring in a tree potentially proxies the rainfall conditions during the period when the ring was formed. A negative correlation consistent with the theoretical expectation was found between rainfall and the *δ*^13^C values of baobab rings in Southern Africa (Woodborne et al., 2015). Low *δ*^13^C values infer higher rainfall while high *δ*^13^C values are indicative lower precipitation, but a direct transfer function accounting for evaporation, rechange and runoff has not been established, and so the proxy record reflects relative changes in effective rainfall through time. The isotopic records from southern African baobabs allow the investigation of the local rainfall drivers of rainfall at a higher spatial resolution, and they reveal decadal, multi-decadal and centennial variability of rainfall over the last 1000 years (Woodborne et al., 2015; 2016). In this paper, we describe new proxy rainfall records for the southwest region of Madagascar during the last millennium based on *δ*^13^C time series from several baobab trees and evaluate the potential drivers of centennial variability of rainfall in this area.

## 2. MATERIAL AND METHODS

### 2.1 Study setting

This research focuses on southwest Madagascar, the driest region of the island (Fig. 1). The area has a semi-arid climate with irregular rainfall (Donque, 1975; Jury, 2016) decreasing from 600 mm to less than 300 mm per year towards the south and the coast. Most of the rainfall occurs during the austral summer (November – March) with monthly rainfall between 300 mm for January and <2 mm in June, July, and August with an above average dry season rainfall recorded between 1983 and 1990, while wet seasons had elevated precipitation 2000 – 2015 (World Bank 2017).

This region supports three baobab species, *Adansonia grandidieri, A. za* and*, A. rubrostipa,* and all three species offer the opportunity to establish paleoclimate records as they are long-lived with distinct radial growth rings (Patrut et al., 2015; Woodborne et al., 2015, 2016).

### 2.2. Sample collection

Sampling was conducted in 2015 on four living trees following a north-south transect of southwest Madagascar to cover the regional climate. The trees were coded as DFL, DFS, GTR and TSP based on their location from north within the dry forest vegetation to the spiny thickets vegetation in the south (shown in Fig. 1; Table 1). The trees were selected and cored, based on their large size (>7 m in circumference) and location (>1 km from any possible surface water source, such as marshes and lakes that could obscure the rainfall signal due to the buffering effects of local hydrology). In addition to official permits for sample collection and exportation from the Ministry of Environment and Sustainable Development of Madagascar and Madagascar National Parks, local communities were consulted and gave permission for the sampling to proceed. Cores were carefully extracted from selected trees using a Haglöf 10mm diameter increment borer at a position of 1.30 m above the ground. After core extraction from each tree, the hole was sealed with a commercial tree sealant to prevent any potential damage to the tree by insects or fungus infestation (Tsen et al., 2015).

**Table 1:** Core information from trees sampled along a north-south transect in southwestern Madagascar.

| Species<br>and<br>sample ID | Coordinates | Elevation (m) | Core length (cm) | Tree circumference at breast height (m) | Number of $\delta^{13}\text{C}$ samples | Number of $^{14}\text{C}$ AMS dates |
| --- | --- | --- | --- | --- | --- | --- |
| *A. gran* 01 – DFL | -20.2224167 °S<br>44.4277500 °E | 25 | 97.5 | 8.8 | 433 | 12 |
| *A. gran* 04 – DFS | -20.8207222 °S<br>44.3940000 °E | 20 | 92 | 11.9 | 445 | 12 |
| *A. gran* 05 – GTR | -21.8602778 °S<br>43.8670556 °E | 23 | 91 | 12.2 | 450 | 13 |
| *A. za* – TSP | -23.8863889 °S<br>44.2583889 °E | 88 | 139 | 7.7 | 706 | 15 |

### 2.3. Tree chronology

The chronologies of the cores were determined from 52 AMS radiocarbon dates: respectively 12 for DFL, 12 for DFS, 13 for GTR and 15 for TSP (Supplementary Information SI1). Samples for radiocarbon dating were selected based on conspicuous shifts in the *δ*^13^C time series. Samples were processed at iThemba LABS in Johannesburg (South Africa). Samples were pre-treated using the acid-base-acid (ABA) method (Hajdas et al., 2017). This treatment consists of the use of acid 0.5M HCl at 60°C, followed by a weak base of 0.1M NaOH at 60°C, to dissolves humic acids and finally a hot acid wash of 0.5M HCl at 60°C, removing any carbonates that precipitated from modern atmospheric carbon dioxide. Following each step, the samples were rinsed until a neutral pH was obtained (Hajdas et al., 2017). The dates were calibrated with the 2020 Southern Hemisphere calibration from the online calibration programs CALIB 14C version 7.10 (Hogg et al., 2020) and the Calibomb program (http://calib.org/CALIBomb/) from Queens University, Belfast, which uses the bomb carbon dataset of Hua et al. (2013).

The age model uses wiggle matching between the trees to constrain possible radiocarbon calibration intercepts, as was used in previously published tree records from southern Africa (Woodborne et al., 2015; 2016). Linear age interpolation of the chronology was used to assign an age to each *δ*^13^C analysis. The approach is a pragmatic attempt to use the climate data as additional apriori constraints on the age of samples. In Bayesian age models, the relative chronology (*x* is known to be older than *y* based on stratigraphy, or in the case of trees, the radial growth) is used as an *apriori* constraint, and these methods provide a means of propagating analytical errors in a series of radiocarbon dates. Our approach notes that within windows of 50-100 years (the kind of error that may be expected in a radiocarbon age model) there are sequences of extreme rainfall and drought events that are common across different trees, as is the atmospheric *δ*^13^C Suess Effect, and these should be presumed to be synchronous because they are driven by large scale synoptic or earth system dynamics. In this way climatology in addition to the radiocarbon dates provide many more apriori constraints on the chronology than the traditional Bayesian models based solely on the radiocarbon dates. None of the existing Bayesian age model packages provide a convenient way to integrate these additional constraints. The argument has a risk of circularity because the chronology is required to generate the climate record, and the climate record is required to refine the chronology, but all attempts to reconcile climate sequences are constrained by the requirement that the age model must fall within the 1-sigma calibration ranges for the radiocarbon dates (with the exception of patent erroneous radiocarbon dates that show clear age inversions in the radial growth of trees, and which should be dismissed as errors, possibly in sample labeling). An implication of this approach is that a date on an individual tree becomes a constraint on the age model of all the trees for that particular time period. The approach that we use yields a chronology that differs very little from Bayesian models but provides a more coherent climate record that more accurately represents the regional rainfall variability through time. The major flaw in the approach is the inability to calculate errors on the age model, although the additional constraints from the climate record, and the demand for coherence in large scale climate events across trees likely yields a tighter chronology than a traditional Bayesian model based only on the within-tree radiocarbon analyses alone. The record obtained from the isotopic values and the age model, was therefore subject to a 21-year-biweight mean analysis which suppresses uncertainties in the age model and also the localised effect of synoptic changes over stochastic weather processes (Woodborne et al., 2015; 2016). The approach prevents any analysis of temporal patterns that are resolved at less than decadal time scales.

### 2.4. Stable carbon isotope analysis

The *δ*^13^C measurements were conducted at the Stable Isotope Laboratory at the University of Pretoria. Each core was mounted on a wood backing panel so that their axial orientation of the baobab core could be seen. The exposed half of the core was sub-sampled from the bark to the pith while being sensitive to the observable direction of the tree growth ring. The remaining half of the core are preserved as an archive. The number of samples per core is provided in Table 1. Each sample was placed in an individually labelled reaction vessel and subject to a soxhlett extraction in a 2:1 toluene/ethanol mixture to eliminate the mobile and soluble components in the wood. Followed by α-cellulose extraction in 17% sodium hydroxide (NaOH) and 10% sodium chlorite (NaClO_2_) at different concentrations to remove the lignins and eliminate the hemicellulose of the wood. The last steps consisted of covering the samples with 1% Hydrochloric acid (HCl) solution for 10 minutes, rinsed and dried overnight at 70°C(Loader et al., 1997; McCarroll & Loader, 2004; Hall et al., 2009). Once dried, aliquots of the α-cellulose were weighed (0.050 - 0.060 mg) using a Mettler Toledo MK5 microbalance and folded into tin capsules before isotopic analysis. The samples were combusted at 1020°C in an elemental analyzer (Flash EA 1110 Series) coupled to a Delta V Plus stable light isotope ratio mass spectrometer, via a ConFlo III system (all equipment supplied by Thermo Fischer, Bremen, Germany). An in-house running standard of Malaysian wood (*Shorea superba*) with an isotopic value of −28.4‰ was used (Hall et al., 2008; 2009; Woodborne et al., 2015; 2016) to calibrate the results. A blank and in-house standard sample which was calibrated against a large number of international standards was run after every 12 samples. The results were reported in *δ*^13^C relative to Vienna-PDB (VPDB) and expressed as per mille according to the formula:

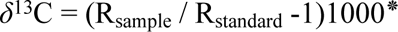

Where *δ*^13^C is the isotopic composition of the sample and R indicates the ratio of ^13^C/^12^C in the sample. Sample replication including sample pre-treatment and error in the analysis were <0.2‰. This is based on a total of 1936 analyses conducted for the overall project, giving 161 standards approximately. The precision achieved here is similar to that reported by other stables light isotope laboratories.

### 2.5. Isotope correction and analysis

Tree physiological responses to environmental conditions are reflected in the *δ*^13^C ratios of the wood tissue (McCarroll & Loader, 2004), but the carbon isotope ratios are also affected by changes in the isotopic ratio of atmospheric CO_2_ over time (Farquhar et al., 1982) and variation in intrinsic water-use efficiency (iWue) in response to elevated atmospheric CO_2_ concentrations since the industrial revolution (Wang & Feng, 2012; Wils et al., 2016). The isotopic data derived from each tree therefore needs to be corrected to isolate the local environmental response. This correction compensates for the *δ*^13^C_atm_ variations using the global record of Belmecheri & Lavergne (2020) and is a simple normalisation to pre-industrial *δ*^13^C_atm_ values (McCarroll & Loader 2004). In this study this is assigned to 1748 (the inflection date between pre- and post-industrial atmospheric changes in the Belmecheri & Lavergne (2020) dataset). In addition to the measured change in *δ*^13^C_atm_, there is also an increase in the atmospheric concentration of CO_2_ (*c*_a_), which affects the intercellular CO_2_ concentration (*c*_i_). Since the rate of carbon assimilation, and accordingly the *δ*^13^C ratio of wood tissue from each growth ring, is linked to the *c_i_/c_a_*ratio (McCarroll & Loader, 2004), reduced water transpiration is associated with elevated *c_a_* during photosynthesis, which increases the iWue of the plant. The correction used in this study followed the method of Woodborne et al. (2016), which is equivalent to the *δ*^13^C pin (“preindustrial”) correction method (McCarroll et al., 2009).

A 21-year biweight mean (a weighted mean that considers the *δ*^13^C values from the decade prior to, and after the date in question) was calculated to emphasize the decadal and multi-decadal pattern in the record. The biweight mean also accommodates the possible age model errors that arise from using a linear growth model to approximate a growth pattern that is likely punctuated by slight variations in growth rates.

The isotopic time series provides a regional decadal record of effective rainfall for the four trees. Each tree experienced unique local climate conditions over the last 700 years; and the composite record allows evaluation of the common and wider climate forcing by emphasizing commonalities in the low frequency component of the record. Change point analysis using the package change point in R was conducted on the composite records. Following this, the composite record was compared with published data for a range of potential rainfall drivers including the local Southwestern Indian Ocean (SWIO) SST, and also equatorial climate shifts caused by the Indian Ocean Dipole (IOD) (Table 2).

**Table 2:** Paleoclimate forcing used for comparison with the baobab isotope records.

| **Data** | **Proxy** | **Time covered**<br>**(AD)** | **Record length**<br>**(years)** | **References** |
| --- | --- | --- | --- | --- |
| Sea Surface Temperature (SST, Ifaty) | Oxygen isotopes of coral | 1660-1994 | 334 | Zinke et al., 2022 |
| Southern Annual Mode Index (SAM) | Mid-latitude to polar domain proxy records | 1000-2007 | 1007 | Abram et al., 2014 |
| Pacific Decadal Oscillation (PDO) | Tree rings | 993-1996 | 1003 | MacDonald & Case 2005 |
| HadISST1.1 SST dataset (Index referring to IOD) | Calculated anomalous SST gradient between the western equatorial Indian Ocean and the south-eastern equatorial Indian Ocean. Based on coral isotopes and Ca/Mg ratios. | 1981-2010 | 29 | Saji & Yamagata 2003 |

The identification of monotonic trends within the record was conducted using a non-parametric Mann-Kendall trend test (Nasri & Modares, 2009; Pohlert, 2018) combined with a Least Squares Regression to evaluate the rate of change in rainfall per year in the regional records. In addition, gridded datasets downloaded from GPCC monthly total precipitation from around Betioky at 2.5° x 2.5° resolution were compared to the composite record. For correlations, significance at a level of 0.05 were accepted.

## 3. RESULTS

The calibrated radiocarbon dates for the four trees suggest that they grew over the last 700 years (1302 – 2013 CE) (Table SI1). The most parsimonious age models, that reconcile the AMS dates and stable carbon isotope records, are shown in Fig. 2. All the trees show linear growth over time except for GTR, which demonstrated a hiatus from 1500 CE to 1700 CE. This is not uncommon in baobabs (Patrut et al., 2017). The age model assigned most of the AMS ages within 1-sigma error of 68%.

**Fig. 2:**
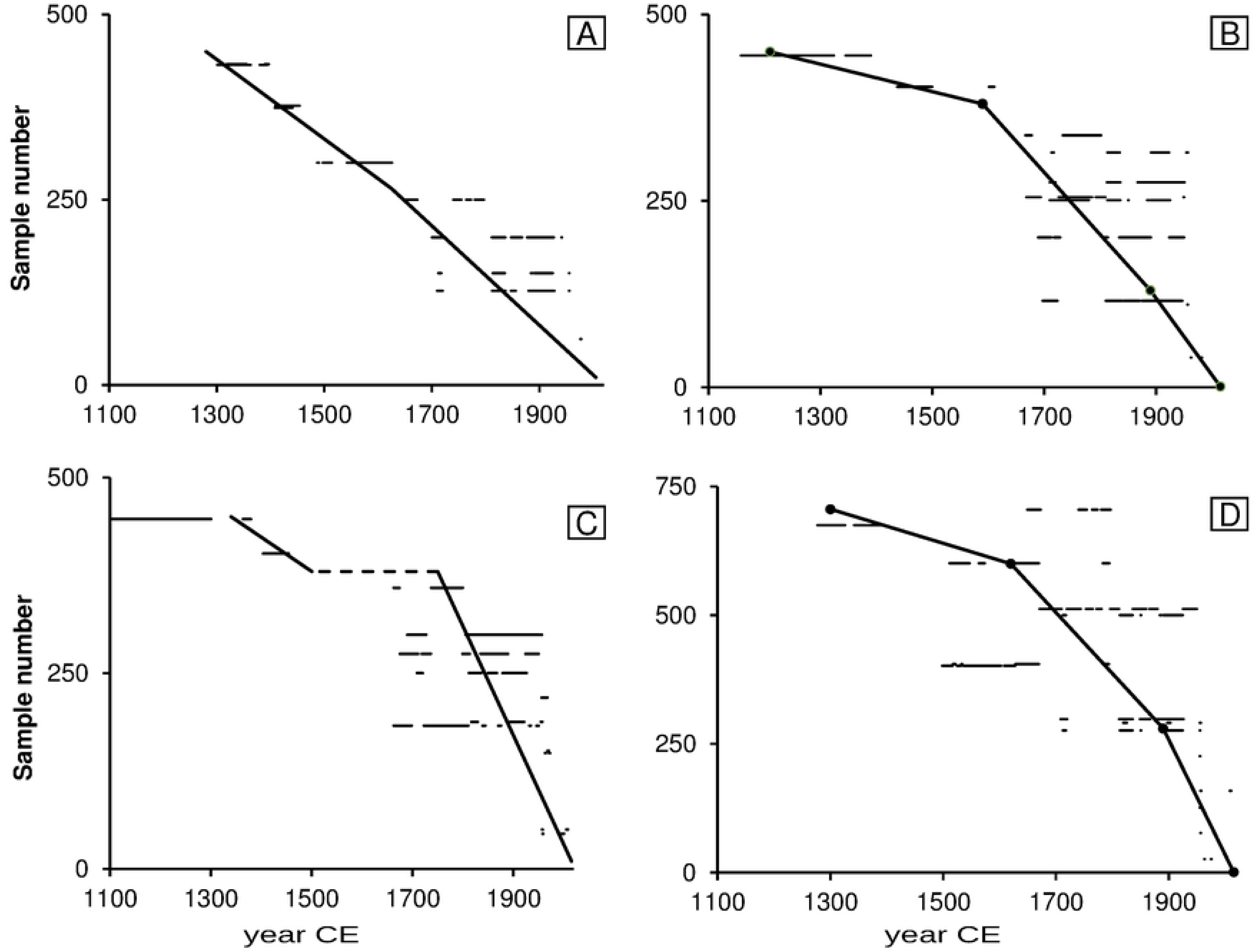
Age-models for (A) DFL, (B) DFS, (C) GTR with the hiatus indicated in dashed line, and (D) TSP, based on 52 radiocarbon dates. The horizontal lines are the 1-sigma calibration intervals for the radiocarbon dates. The bold line represents the age model that best intercepts the 1-sigma calibration range for the radiocarbon dates

The corrected *δ*^13^C time series from the four trees range between −26.2‰ (1400 CE) and −24.5‰ (1650 CE) with a mean of −25.3‰, and a variation of about 1.7‰ (Fig. 3). The trend analysis on the composite *δ*^13^C records using the Mann-Kendall test shows that there is a marginal decreasing trend of the isotope data over time (S=-5.07, p<0.01) with a difference of about - 0.7‰ between 1300 CE and 2013 CE. A least square linear regression is significant (F= 80.6, p = 0.001) but with an R^2^ of 0.10. The change point analysis revealed a number of major shifts in the mean values of the composite isotope data becoming either more positive or negative in the time series (significant at 95%). These occur at approximately 1400 CE, 1480 CE, 1500 CE, 1630 CE, 1660 CE, 1820 CE, and 980 CE (Fig. 4).

**Fig. 3:**
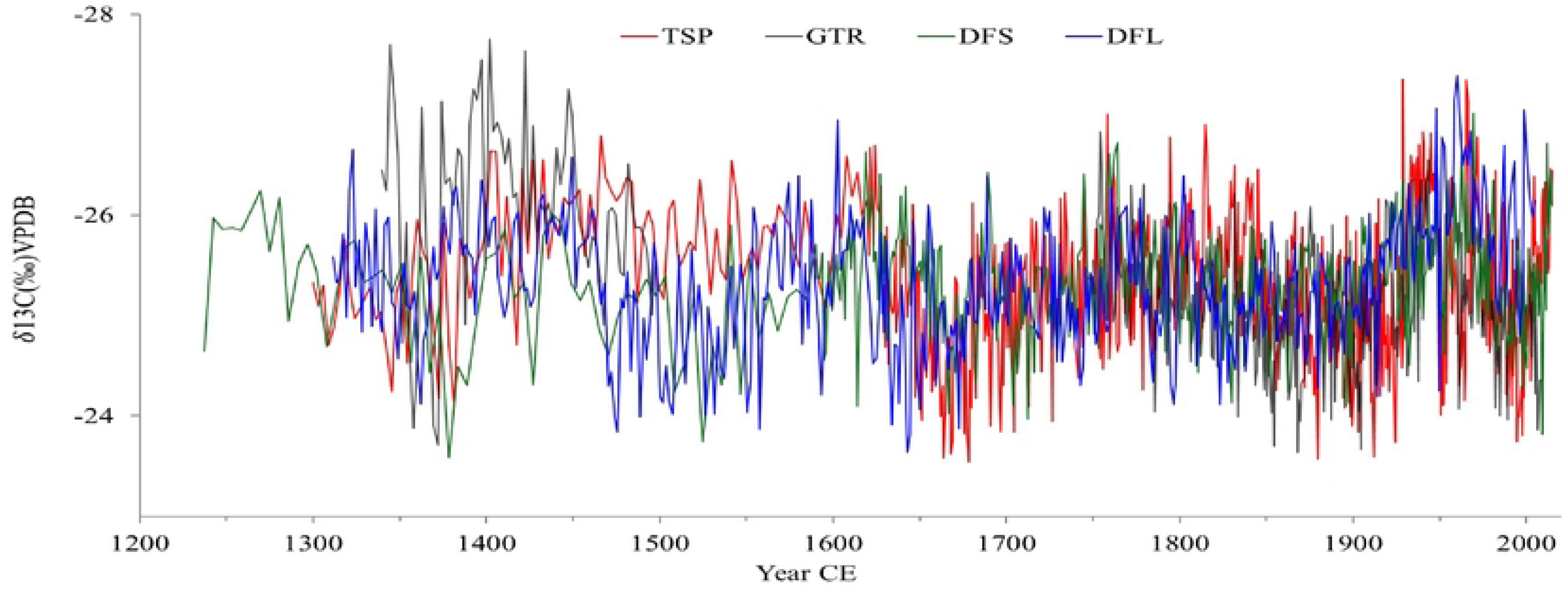
Corrected *δ*^13^C time series of four baobabs from southwest Madagascar with inverted y-axis indicating drier (less negative isotope value) and wetter conditions (more negative isotope value).

**Fig. 4:**
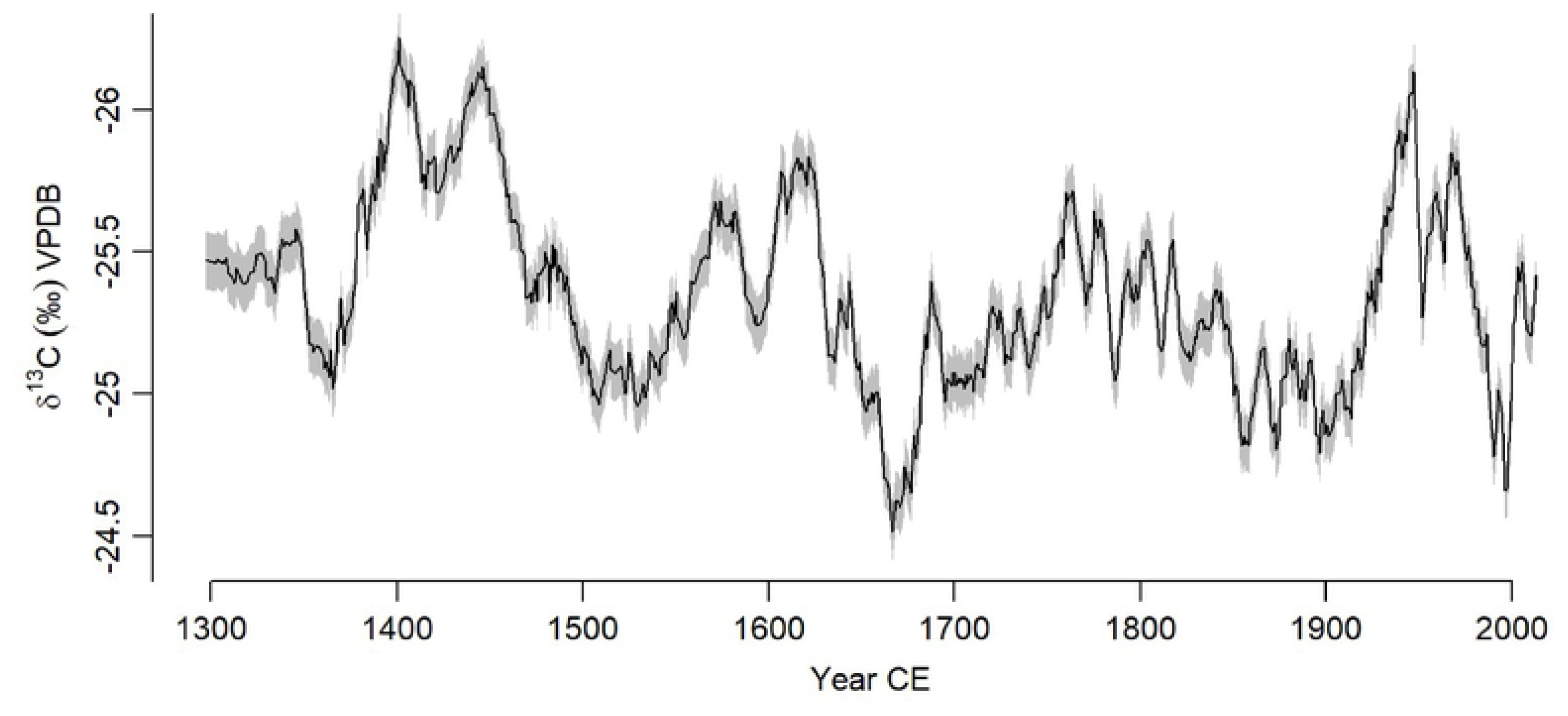
The composite record from four trees (black) with wetter periods (blue) and drier periods (yellow). Error bars represent standard errors.

The isotope biweight mean composite and the GPCC monthly total precipitation both shows similar trends over time. A linear regression of precipitation data from the GPCC monthly total precipitation from 1900–present revealed a decreasing but not significant trend over time (β = - 0.67, p = 0.17), with year explaining only ∼1.7% of the variance (R² = 0.017). Conversely, the isotope data showed a small but significant increasing trend (β = +0.00112, p = 0.015) reflecting decreasing rainfall, though the model explains less than 1% of variance (R² = 0.0099). These results suggest subtle but differing temporal behaviours in the two proxies potentially associated with the associated ages. Both records show a slight decrease in rainfall as reflected by the precipitation value and the more positive isotope value around 1950 and then from 1970-1990 while more wetter periods are recorded around 1960 and 2000 (Fig. 5).

**Fig. 5:**
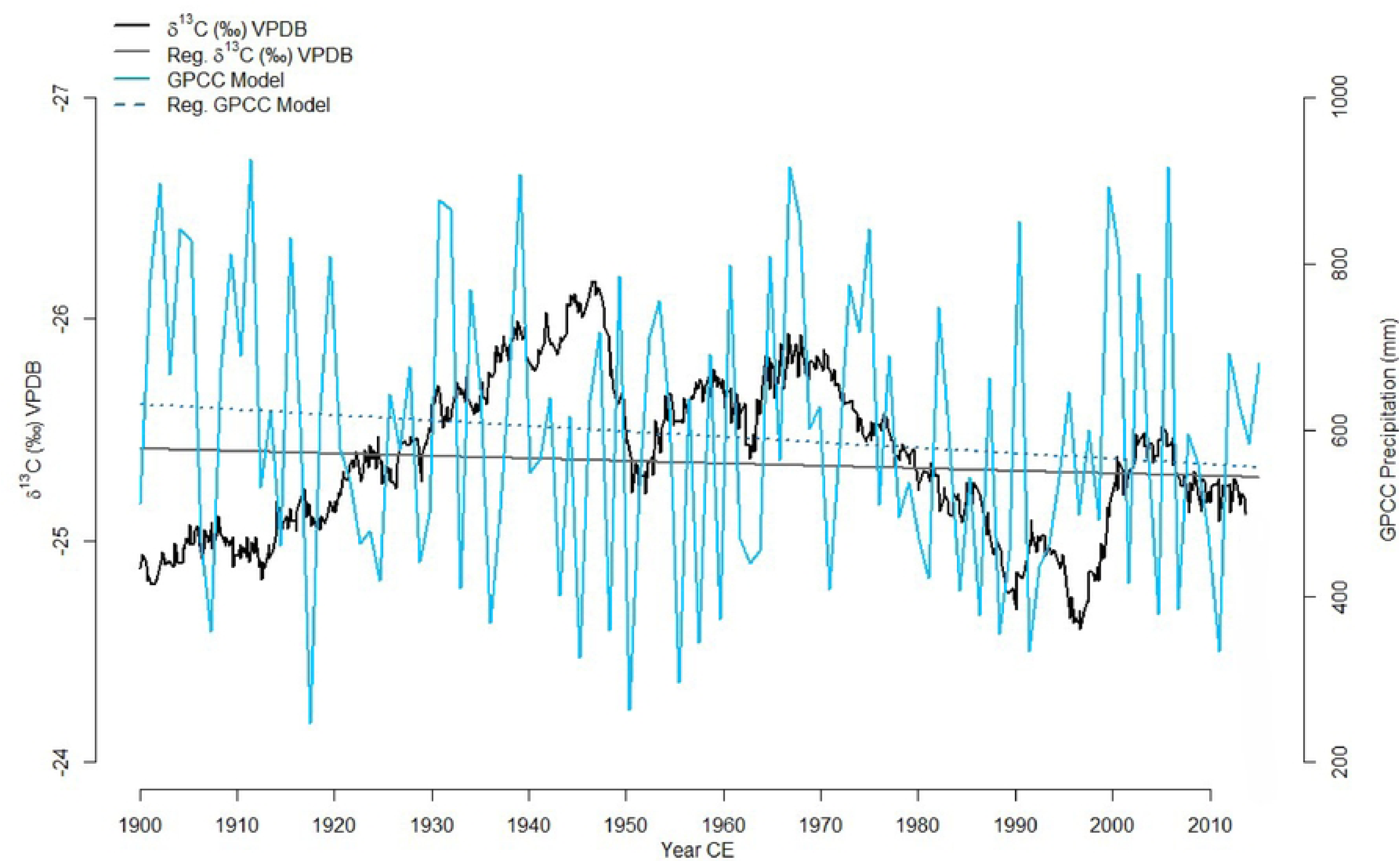
Comparison of the baobab *δ*^13^C records with existing model datasets: Black indicates the baobab *δ*^13^C composite records from 1900-2015 with linear regression indicated in grey. Blue and dark blue shows GPCC total precipitation at 2.5° x 2.5° resolution from around Betioky between 1900-2013 along with the associated linear model.

The correlation analysis of potential rainfall drivers in Southwest Madagascar has been calculated (Table 3). The results diverse correlation values and significance (Table 3). There is a negative correlation between SST and the isotope data an overall significant positive correlation between SAM and the isotope records with recorded weak correlation since the 16^th^ century.

**Table 3:** Correlation between baobab isotope data and various environmental indices. Pearson correlation coefficients (r) and associated p-values are shown. Sample size (n) indicates the number of data points compared for each dataset.

| Dataset Compared with Baobab Isotope Data | Time Period (AD) | Sample Size (n) | Pearson Correlation (r) | p-value |
| --- | --- | --- | --- | --- |
| Sea Surface Temperature (SST, Ifaty) | 1660–1994 | 334 | −0.22* | < 0.001* |
| Southern Annual Mode Index (SAM) | 1000–2007 | 1007 | 0.16* | < 0.001* |
| Pacific Decadal Oscillation (PDO) | 993–1996 | 1003 | 0.01 | > 0.05 |
| HadISST1.1 SST dataset (IOD Index) | 1981–2010 | 29 | −0.06 | > 0.05 |

Note: \* indicates significance at $\alpha = 0.05$ .

Related to the Pacific Decadal Oscillation (PDO) reconstruction, no significant correlation has been recorded during the entire period but during the LIA negative PDO anomalies are associated with decreases in rainfall particularly between 1600 – 1750 CE (r=-0.30, p<0.001), whereas in the record before and after this period, they are associated with increases in rainfall (r=0.18, p=0.004 and r=0.18, p=0.001 respectively) (Fig. 7C).

## 4. DISCUSSION

### 4.1. Rainfall record of southwest Madagascar for the last 700 years

The similarities between the *δ*^13^C records and the GPCC precipitation data meets the theoretical expectation, and reaffirms the results obtained for baobab isotope controls on the African mainland (Woodborne et al., 2015; 2016; Fig. 5). The baobab *δ*^13^C record can thus be interpreted as a proxy for local effective rainfall in southwest Madagascar, reflecting decadal to centennial variability in the last 700 years. Accordingly, lower *δ*^13^C values indicate wetter periods while more positive isotope ratios correlate to drier conditions.

The chronology of the four baobabs ranges from 1300 CE until 2013 with a hiatus in the GTR core from 1500 CE to 1700 CE. Punctuated growth models have been noted in other baobabs (Patrut et al., 2017) and may arise because of attenuated growth rates over time, or possibly due to lobate development of the stem resulting in the reallocation of resources to another part of the tree. The GTR baobab was collected near the River Mangoky in southwest Madagascar where a sediment core was taken from nearby Lake Tsizavatsy. The age-depth model of the Tsizavatsy core shows a hiatus from 1400 to 1900, suggested to be associated with a regional drought event (Razanatsoa et al., 2021). Although the other trees show positive isotope excursions (dry events) during this period, there are oscillations of wetter conditions covered by the present records (Fig. 3). Notwithstanding the gap, the record from GTR prior to and after the hiatus was combined with the composite record of all trees to provide a full rainfall proxy record for the last 700 years for southwest Madagascar covering the Little Ice Age (LIA; Lechleitner et al., 2017; Putnam & Broecker, 2017) and the Anthropocene.

Synchronicity in the baobab *δ*^13^C records from southwest Madagascar demonstrate regional variability with a succession of wet and dry cycles on the scale of decades to centuries (Fig. 4). The rainfall in the region was suggested to be variable with drought events recorded in Southwest Madagascar prior 1000 CE based on stalagmite records (Faina et al., 2021) with drying trends being recorded on the isotope tree records since 1300 CE. Both records, baobab δ^13^C and stalagmite δ¹⁸O show similar trends and patterns demonstrating high agreement in the variability of rainfall over time despite a weak negative Pearson correlation between the two variables (See Table 3). This non correlation could be explained by the uncertainties associated with the two age models. Before 1700 CE, both records show different trends with discrepancies around mid-17^th^ century where the tree record showed drier conditions while the stalagmites is showing wetter conditions. Post 1700 CE similarities between the stalagmite *δ*^18^O and *δ*^13^C tree records are recorded with wet condition around 1700-1800 CE, dry condition 1800-1900 CE and a wetter condition again between 1900-2000 CE. Very recently (post-1950 CE), both records reflected a severe drought (Fig. 6).

**Fig. 6:**
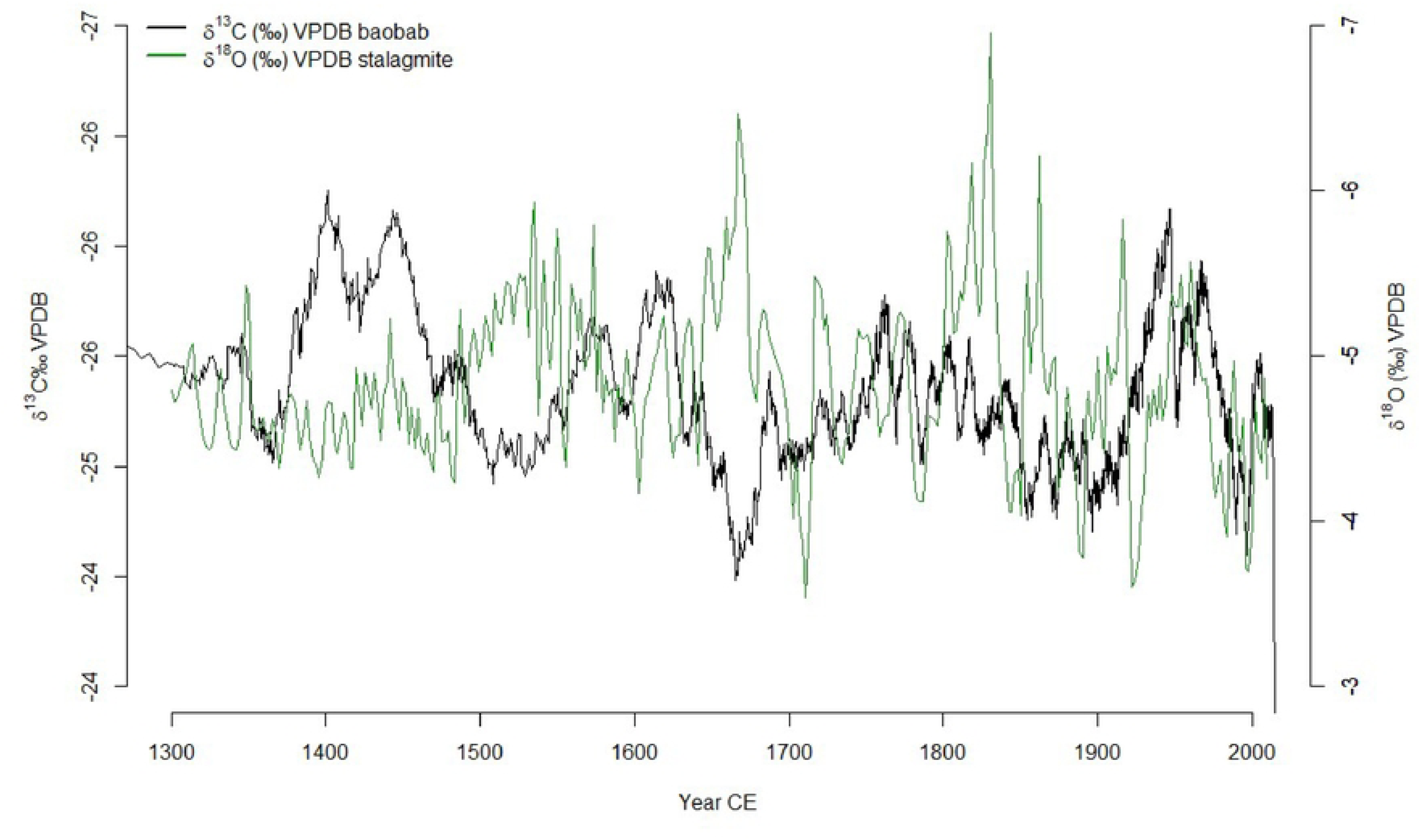
Comparison between baobab *δ*^13^C records from tree rings in southwest Madagascar (black) with stalagmite *δ*^18^O records from Asafora cave from the southwest coast and just southeast of the Velondriake Marine Reserve (green, from Faina et al., 2021).

The composite baobab record shows increasing *δ*^13^C values indicating marginal drying at the locations of the trees over the past 700 years, corresponding with previous findings suggesting increasing aridity since 1000 CE (e.g. Burney, 1993; Burns et al., 2016; Virah-Sawmy et al., 2016, Razanatsoa et al., 2021). Pollen records of the last millennia show a synchronous desiccation from various regions in Madagascar including the southwest (Burney, 1993; Burns et al., 2016; Virah-Sawmy et al., 2016) which agrees with reduced length of wet periods in the tree records compared to regional records.

### 4.2. Synoptic drivers of rainfall in southwest Madagascar

When compared with other baobab *δ*^13^C records from South Africa (Woodborne et al., 2015; 2016), the rainfall patterns for southwest Madagascar and the South African summer rainfall zone are in phase for most of the last 700 years. Wet (dry) periods in the South African Pafuri and Mapungubwe records (22 °S) corresponded to similar wet (dry) periods from southwest Madagascar. This suggests that southwest Madagascar rainfall responds to similar drivers of the summer rainfall zone in southern Africa, including Agulhas Current sea-surface temperature variations regulated by the East-West displacement of the TTTs (Woodborne et al., 2016) in addition to the regulatory effect of the island’s mountains (Donque, 1975; Jury & Huang, 2004). However, further investigation of the various drivers of rainfall is required to provide more understanding of local and regional temporal changes.

#### 4.2.1. ITCZ and SAM modulated rainfall during Early Little Ice Age between 1370 and 1500

The composite baobab record from southwest Madagascar shows a wet period between approximately 1370 – 1500 CE (Fig. 4) which coincides with the second phase 1495 – 1833 CE of a wet-neutral-wet cycle recorded in northwest Madagascar (Scroxton et al., 2017). The movement of the ITCZ has a significant impact on rainfall variability in Madagascar (Haug et al., 2001; Liu et al., 2003; Verschuren et al., 2000; Schneider et al., 2014). A southward shift was recorded at the beginning of the Little Ice Age around 1300 CE (Chiang & Bitz, 2005; Broccoli et al., 2006; Lechleitner et al., 2017; Putnam & Broecker, 2017) and this may have resulted in increased rainfall for southwestern Madagascar. Wetter conditions are also evident in several East African sediment records during this period, including Lake Chilwa (Crossley et al., 1984), Lake Malawi (Johnson et al., 2001), Lake Massoko (Barker et al., 2000), Lake Tanganyika (Alin & Cohen 2003), Lake Victoria (Stager et al., 2005), Lake Naivasha (Verschuren et al., 2000; Tyson et al., 2001; Tierney et al., 2013) suggesting a common driver of rainfall, most likely the southwards movement of the ITCZ.

The wetter conditions from 1370 – 1500 CE in southwestern Madagascar occur when the Southern Annular Mode (SAM) was at its most extreme negative phase in the fifteenth century (Abram et al., 2014; Fig. 7A). Negative SAM indices imply an expansion of the westerly circumpolar vortex, which has significant impacts on temperature and precipitation over all four Southern Hemisphere continents (Gillett et al., 2006), and particularly over Africa south of 25°S, where an increase in precipitation is associated with the northward migration of the westerlies during the austral winter. The response is most pronounced along the east coast, where it is associated with anomalous easterly winds advecting more moisture off the SWIO (Gillett et al., 2006). No significant response to SAM has been reported for Madagascar on the basis of instrumental data (Gillett et al., 2006) although heavy rainfall events have been associated to atmospheric circulation displays a Southern Annular Mode-like pattern throughout the hemisphere (Randriamahefasoa & Reason, 2017). The relationship between SAM and rainfall in the baobab record presented here is not consistent with an overall significant positive correlation (r=0.16, p<0.001). However, since the 16^th^ century the relationship is weak and non-significant suggesting that the southward contraction of the westerly winds during the positive SAM reduces their influence on the region. When the westerlies migrate southward during a positive SAM phase, they cease to be a driver of rainfall, and decadal to centennial rainfall variability responds to other forcing. The lack of consistent correlation throughout the records also supports recent findings related to the lack of a forced response in SAM variability prior to the 20th century (King et al 2023).

**Fig. 7:**
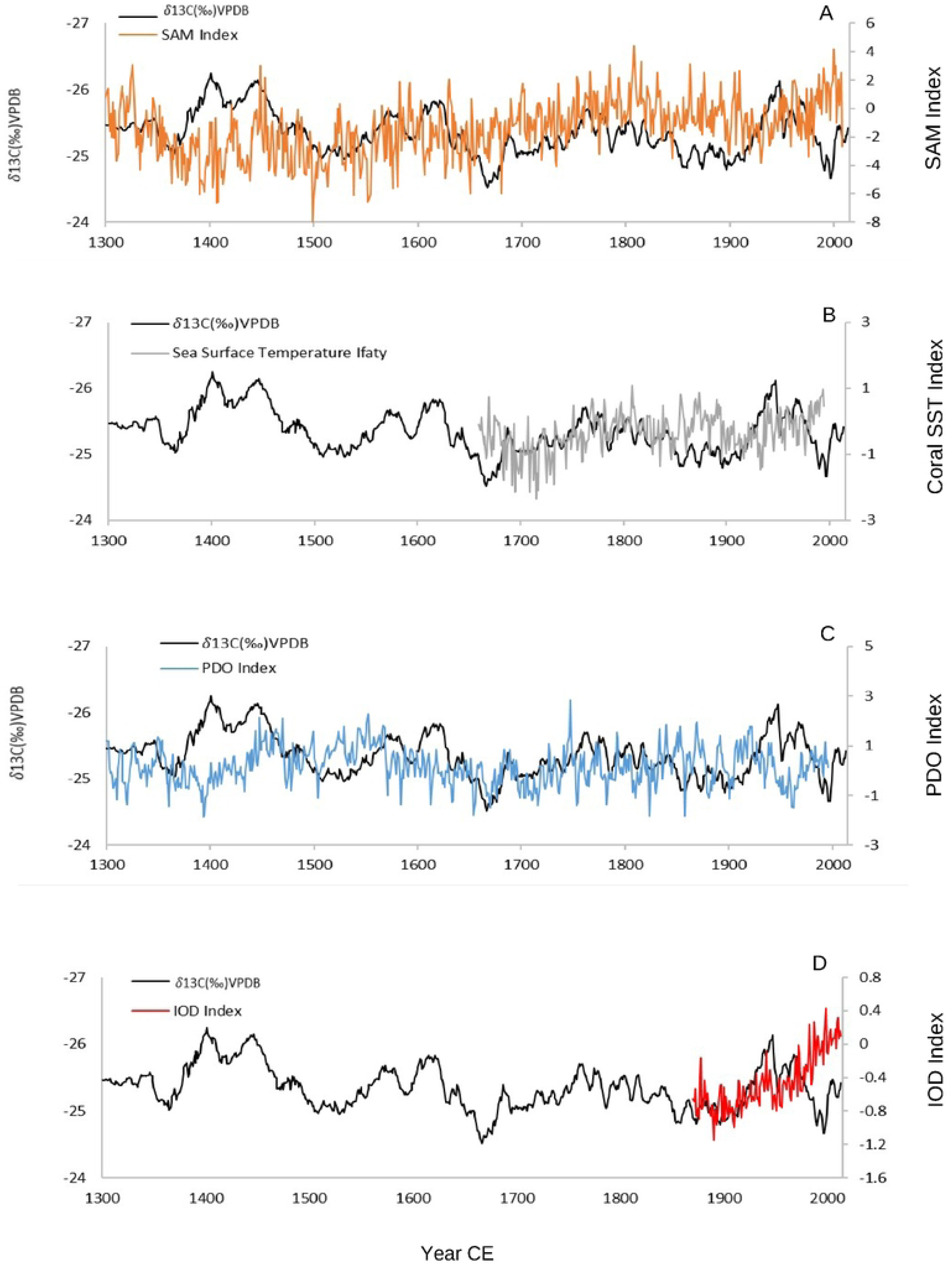
Comparison of the baobab *δ*^13^C records (black) with (A) the Southern Annual Mode (SAM) index (orange) (Abram et al. 2014), (B) the Pacific Decadal Oscillation (PDO) index (**blue**) (MacDonald & Case 2005), (C) Sea Surface Temperature (SST) from Ifaty, southwest Madagascar (gre**y**) (Zinke et al. 2022), and (D) HadISST1.1 SST dataset that refers to Indian Ocean Dipole variability **(red, Saji & Yamagata 2003**)

Our composite record suggests that rainfall responds to subtropical forcing during extreme negative SAM phases as the westerly winds migrate northwards bringing wetter condition to the subtropics including southern Madagascar (Fig. 7A), but otherwise it responds to tropical forcing determined by the position of the ITCZ during the austral summers. Evidence of this mechanism operating during glacial periods is derived from marine records and model analysis of the late Glacial maximum over southern Africa (Anderson et al., 2009; Sigman et al., 2010; Miller et al., 2019b; Engelbrecht et al., 2019; Hahn et al., 2021). The mechanism may explain the rainfall maximum during the 14^th^ and 15^th^ centuries, but further research is needed to elucidate the influence of the SAM on southern African climate during the Little Ice Age.

#### 4.2.2. Regional and localised rainfall drivers during the LIA 1600-1750

Wet conditions during the 14^th^ and 15^th^ centuries are followed by extremely dry conditions 1600 – 1750 CE in southwest Madagascar. This period is characterised by a positive phase of the SAM that commences at 1500 CE (Abram et al., 2014) implying reduced effect of the westerlies. It is also a period of reduced sunspot activity known as the Maunder Minimum (1645 – 1715) (Mann et al., 2009; Scroxton et al., 2017) with decreased global temperatures at the maximum of the Little Ice Age (LIA; Tyson et al., 2001). It has been speculated that there was a southward migration of the ITCZ during the maximum of the LIA (Jury & Huang, 2004; Russel et al., 2007; Tadross et al., 2008; Voarintsoa et al., 2017). Our results show drier condition during this period starting around 1600 CE and has also been experienced in northwest of Madagascar around 1700 CE (Scroxton et al., 2017). Dry periods were also experienced across the African continents including East Africa (Russell & Johnson, 2007; Tierney et al., 2013), and the southern African summer rainfall area (Huffman, 2004; PAGES 2k Consortium, 2013; Macron et al., 2014; Chevalier & Chase, 2015; Huffman & Woodborne, 2016; Woodborne et al., 2016). Pollen evidence from the Lake Longiza in southwest Madagascar suggested an increase in grass and decrease in trees and shrubs such as Arecaceae, *Pandanus*, and *Acacia* potentially associated with the drying in the region (Matsumoto & Burney, 1994; Razanatsoa et al., 2022). Lake Tsizavatsy, from the same region, showed a hiatus in its sediment deposition between 1400 – 1900 CE (Razanatsoa et al., 2021) suggesting the drying of the lake during this period, which is consistent with the tree records. The evidence of dry conditions from records across southern Africa does not support southward migration of the ITCZ during this period and if the migration occurred, there might have been other drivers that inhibited its effect leading to a decrease in rainfall during this period.

The Pacific Decadal Oscillation (PDO) is linked to ENSO in relation to drought patterns across Africa through teleconnections that influence the longitudinal position of climate systems. The (PDO) reconstruction shows that during the LIA negative PDO anomalies are associated with decreases in rainfall particularly between 1600 – 1750 CE (r=-0.30, p<0.001), whereas in the record before and after this period, they are associated with increases in rainfall (r=0.18, p=0.004 and r=0.18, p=0.001 respectively) (Fig. 7C). Despite the change in the sign of the correlation, this evidence suggests that climate forcing is dominated by tropical forcing. (Thompson et al., 2003; MacDonald & Case 2005; Hoell et al., 2017). Changes in the TTTs position which tend to propagate eastward, from southern Africa to the Mozambique Channel and southern Madagascar is known to have a strong influence on intra-seasonal and even interannual rainfall variability in the region (Macron et al., 2014). This has particularly been suggested to be dictated by the migration of the ITCZ (Chase & Meadows, 2007; Sachs et al., 2009) and to increase during La Nina conditions (Manhique et al., 2011; Ratna et al., 2012; Macron et al., 2014). Moreover, its persistence was suggested to be maintained by variation of SST anomalies over the Agulhas Current (Manhique et al., 2011; Vigaud et al., 2007). Comparison of the baobab *δ*^13^C records from 1660 – 1994 CE with the 300-year Agulhas Current Sea surface temperature (SST) record from Ifaty in southwest Madagascar (Zinke et al., 2014) shows a significant negative correlation (r=-0.22, p<0.001). Positive (negative) SST corresponds with higher (lower) rainfall in the baobab record (Fig. 7B). The coolest oceanic temperatures in the coral record, with anomalies of −0.3 – −0.5 °C between 1675 – 1760 CE, correspond to the driest period in southwest Madagascar. Similar patterns of rainfall were noted in the summer rainfall area of the adjacent African mainland (Woodborne et al., 2015). Variation in SST in the western Indian Ocean are determinant in the IOD with a suggested negative relationship with eastern African rainfall responses (Hoell et al., 2017; Taylor et al., 2021).

#### 4.2.3. Mixed effect of changes in ITCZ position, human land-use and climate change from 1750 – 2013 CE

Around 1750 CE until early 1800 CE, at the end of the LIA, there is a relatively wet period recorded in the baobab *δ*^13^C data. A relatively dry period after 1860 CE is similar to conditions experienced over the summer rainfall zone in southern Africa. These periods coincided with more extreme ENSO warm phases (Nash, 2017) with the warmest period in the Agulhas SST record between 1880 CE and 1900 CE and a northward migration of the ITCZ (Zinke et al., 2014; Railsback et al., 2018).

The comparison of the composite baobab *δ*^13^C record with the Dipole Mode Index (DMI) that were used as an index of the IOD (1870 – 2013) shows very low to no correlation (Table 3, Fig. 7D) but with a noticeable positive but not significant correlation since 1980 CE (r= 0.27, p= 0.08). The effect that the IOD has on equatorial climate forcing is similar to the ENSO or PDO effects as it is driven by an equatorial SST differential across the Indian Ocean while ENSO is driven by a gradient in the Pacific Ocean. The effect in the subtropical region of southwestern Madagascar appears to be dominated by the Pacific Ocean influences on global climate (ENSO/PDO) which is influential in the latitudinal position of the TTTs system.

Around 1950 CE, conditions in southwest Madagascar are as wet as at any time in the record. This corresponds to a positive SST anomaly, positive phase of IOD and a negative PDO phase. Despite suggested changes in the impact of ENSO cycles on the SST in the region of the SWIO since 1970 (Zinke et al., 2014), our results show typical PDO/rainfall phasing with more negative PDO corresponding to high rainfall while the inverse is not always true. Records suggest that the IOD intensified following the onset of global warming during the 20^th^ century along with forced response of SAM (Abram et al., 2008; Namakura et al., 2009; Watanabe et al., 2019; King et al 2023), with increased evidence of human induced climate change (IPCC, 2021), evidence of increased river runoff and shifts in human land-use through slash and burn agriculture were recorded in coral records from eastern Madagascar (Grove *et al.,* 2013). Pollen records from the region suggest a decrease in the tree component including Arecaceae coinciding with an increase in Poaceae and pioneer taxa such as Asteraceae mostly likely associated with tree cutting associated with agriculture expansion (Razanatsoa et al., 2022). These suggest that changes in rainfall in the region led to changes in land-use. There is a return to drier conditions around 1980 called “belt of iron” also recorded in historical records peaking at the beginning of the 1990s (Von Heland & Folke, 2014). This was followed by a trend towards wet conditions in the past 20 years similar the instrumental record (Tadross et al., 2008).

### 4.3. Implications for future climate change risk and adaptation

The last 300 years of the baobab composite record show the interacting global, regional and local drivers that have influenced rainfall variability in southwest Madagascar. The dominant effect of the position of the ITCZ during the austral summer is evident, and this is troubling in the context of forecast climate change. The position and zone of the ITCZ is predicted to narrow with a northward shift over eastern Africa and the Indian Ocean and a southward shift in the eastern Pacific and Atlantic oceans by 2100, which would severely reduce rainfall in the region (Mamalakis et al., 2021). In terms of climate risk and adaptation, southwest Madagascar will likely experience a drier climate with more frequent and prolonged droughts as already predicted in the IPCC 6^th^ assessment report (IPCC, 2021). Climatic factors are an important driver of economic, environmental and societal decisions (Boutin & Smit, 2016) and would be crucial for the future of these dry areas. Indeed, the population are dependent on rainfall as a source of water and for agriculture due to the lack of infrastructure and the limited permanent water ponds (Hänke et al., 2017; Carriere et al., 2018). The effect of drought and lack of rainfall has already led the Mikea forager communities to diversify their livelihoods with seasonal agriculture to ensure food security (Razanatsoa et al., 2021). Some adaptations that have been established elsewhere and could be conducted in the region include the introduction of new drought resistant crops, (e.g. Thomas et al., 2007; Yaro et al., 2014). Multiple species livestock herding with cattle and goat pastoralism (Kaufmann & Tsirahamba, 2006; Hänke & Barkmann, 2017) and livelihood diversification including work that is not farming (e.g. crafts and services for local markets) were suggested to be a major adaptive strategy under drying conditions in a short and long term and buffer livelihoods in the face of environmental change (e.g. Kuiper et al., 2007; Tambo & Abdoulaye, 2013). Further understanding of the effect of future climate on these populations and their surrounding environment is critical in planning strategies of adaptation in terms of livelihoods but also water provision also in the coming years.

## 5. CONCLUSION

Baobab *δ*^13^C data from southwest Madagascar are a proxy for changing rainfall over the last 700 years. The inferred wettest period was between 1370 – 1500 CE while the driest period occurred between 1600 – 1750 CE. High centennial variability was recorded with a decreasing rainfall trend and reducing duration of wet periods over time. The comparison of the records with existing records of rainfall drivers at local, regional, and global scales shows that the baobab rainfall proxy record is not dominated by the influence of any single forcing over the entire record. The westerlies may play a role during extreme negative phases of the SAM, while latitudinal shifts of the ITCZ are the dominant low frequency driver of rainfall. At a more local level, the role of SST seems to dominate variability possibly through longitudinal influences on the position of the TTTs system which is also influenced by the PDO/ENSO system. Localised climate forcing in relation to the Southwest Madagascar Coastal Current (SMACC) within the greater Agulhas Current system has been suggested (Ramanantsoa et al., 2018). What emerges from comparisons with other rainfall proxy records on the island of Madagascar, and from the adjacent African mainland, is that the temporal trends are not consistent, probably reflecting the contrasting dominance of different drivers in different regions. A rainfall dipole exists between southern Africa and Madagascar (Jury et al., 2015; 2016; Woodborne et al., 2016; Barimalala et al., 2018) with increases in precipitation over southern Africa extending from Mozambique to Angola coincident with a decrease in rainfall most of Madagascar (Barimalala *et al.,* 2018) but not in the southwest region. The mountains that extend from the north to the south of Madagascar (>1500 m elevation) reduce the direct transport of moisture from the Indian Ocean toward southern Africa (Barimalala et al., 2018) and southwest Madagascar. The Agulhas current SST forcing of Madagascar rainfall is opposite to that in southern African where negative SST anomalies were associated with wetter conditions over the southern African interior (Woodborne et al., 2015). These contradictions suggest that the variation in rainfall is not a simple intensification or weakening of the existing climate patterns, but rather a response in the synoptic systems to tropical (ENSO/PDO), extratropical (SAM), and localised (SST) forcing. Why some forcing appears to dominate at certain periods and not at others is unclear, and the evidence presented here suggests that synergistic effects might be explored in global climate models.

The potential effect of climate change and land-use change were also recorded at the near present period, as well as possible effects of SAM if there is a northward migration of westerlies similar to what happened around 1300 CE. The data generated here provide the opportunity to unravel the relative importance and interaction between global, regional, and local drivers across the southern and eastern African region. These findings are crucial in the simulations of rainfall projections to help evaluate the impacts and trends of migration of the westerlies and anthropogenic induced climate change on the African continents that are not fully understood. For southwest Madagascar with an expected drier climate and increasing occurrence of severe drought conditions predicted for the near future, understanding the risks, and establishing adaptation strategies particularly in terms of livelihood could avoid disastrous famine.

## 1. ACKNOWLEDGEMENT

We would like to thank the Ministry of Environment and Sustainable Development of Madagascar and Madagascar National Park for providing the permission for the field campaign, sampling and exportation. We acknowledge ESSA-Forêts Mention Foresterie et Environnement de l’Ecole Supérieure des Sciences Agronomiques, Université d’Antananarivo – MADAGASCAR for collaborating with the obtention of the research permit. We also acknowledge all the field assistants that have participated in retrieving the cores and Tsilavo Razafimanantsoa for his input on the map.

## 2. DATA AVAILABILITY

Datasets related to this article can be found at 10.25375/uct.16590035 on Zivahub, an open-source online data repository hosted by the University of Cape Town. The DOI will become active upon publication of the manuscript.

## Supplementary Information

**SI1:** Radiocarbon dates of each tree replicate from the four trees collected in southwest Madagascar

## DATA REFERENCES

Zinke, J., B. Loveday, C. Reason, W.-C. Dullo, and D. Kroon. (2014). Madagascar corals track sea surface temperature variability in the Agulhas Current core region over the past 334 years. Scientific Reports, 4, 4393. doi: 10.1038/srep04393. NOAA’s National Centers for Environmental Information (NCEI)

Abram, N. J., Mulvaney, R., Vimeux, F., Phipps, S. J., Turner, J., & England, M. H. (2014). Evolution of the Southern Annular Mode during the past millennium. Nature Climate Change, 4(7), 564–569. 10.1038/nclimate2235. NOAA’s National Centers for Environmental Information (NCEI)

Jinbao Li, Shang-Ping Xie, Edward R. Cook, Gang Huang, Rosanne D’Arrigo, Fei Liu, Jian Ma, and Xiao-Tong Zheng. 2011. Interdecadal modulation of El Niño amplitude during the past millennium. Nature Climate Change. 1(2) 114–118. doi: 10.1038/nclimate1086. NOAA’s National Centers for Environmental Information (NCEI)

Scroxton, N., Burns, S. J., Mcgee, D., Hardt, B., Godfrey, L. R., Ranivoharimanana, L., & Faina, P. (2017). Hemispherically in-phase precipitation variability over the last 1700 years in a Madagascar speleothem record. 164. 10.1016/j.quascirev.2017.03.017. National Centers for Environmental Information (NCEI)

MacDonald & Case. 2005. Pacific Decadal Oscillation Reconstruction for the Past Millennium. GRL 32, L08703. doi:10.1029/2005GL022478. National Centers for Environmental Information (NCEI)

Saji, N.H., & Yamagata, T., (2003). Possible impacts of Indian Ocean Dipole mode events on global climate. CLIMATE RES, 25 (2): 151–169. https://psl.noaa.gov/gcos_wgsp/Timeseries/DMI/

## REFERENCES

Abram, N. J., Gagan, M. K., Cole, J. E., Hantoro, W. S. & Mudelsee, M. (2008). Recent intensification of tropical climate variability in the Indian Ocean. Nature Geoscience 1, 849–853. 10.1038/ngeo357

Abram, N. J., Mulvaney, R., Vimeux, F., Phipps, S. J., Turner, J., & England, M. H. (2014). Evolution of the Southern Annular Mode during the past millennium. Nature Climate Change, 4(7), 564–569. 10.1038/nclimate2235

Alin, S. R., & Cohen, A. S. (2003). Lake-level history of Lake Tanganyika, East Africa, for the past 2500 years based on ostracode-inferred water-depth reconstruction. Palaeogeography, Palaeoclimatology, Palaeoecology, 199(1–2), 31–49. 10.1016/S0031-0182(03)00484-X

Anderson, R. F., Ali, S., Bradtmiller, L. I., Nielsen, S. H. H., Fleisher, M. Q., Anderson, B. E., and Burckle, L. H. (2009). Wind-Driven Upwelling in the Southern Ocean and the Deglacial Rise in Atmospheric CO2, Science. 323, 1443–1448, 10.1126/science.1167441.

Baptista, D. M. S., Farid, M., Fayad, D., Kemoe, L., Lanci, L. S., Mitra, P., Muehlschlegel, T. S., Okou, C., Spray, J. A., Tuitoek, K., & Unsal, F. D. (2022). Climate change and chronic food insecurity in sub-Saharan Africa. International Monetary Fund Library, 2022(16), 1. 10.5089/9798400218507.087

Barimalala, R., Desbiolles, F., Blamey, R. C., & Reason, C. (2018). Madagascar Influence on the South Indian Ocean Convergence Zone, the Mozambique Channel Trough and Southern African Rainfall. Geophysical Research Letters, 45(20), 11,380-11,389. 10.1029/2018GL079964

Barimalala, R., Raholijao, N., Pokam, W., & Reason, C. J. C. (2021). Potential impacts of 1.5 °C, 2 °C global warming levels on temperature and rainfall over Madagascar. Environmental Research Letters, 16(4). 10.1088/1748-9326/abeb34

Barker, P., Telford, R., Merdaci, O., Williamson, D., Taieb, M., Vincens, A., & Gibert, E. (2000). The sensitivity of a Tanzanian crater lake to catastrophic tephra input and four millennia of climate change. Holocene, 10(3), 303–310. 10.1191/095968300672848582

Baudoin, M.A., Vogel, C., Nortje, K., Naik, M. (2017). Living with drought in South Africa: lessons learnt from the recent El Niño drought period. International Journal of Disaster Risk Reduction., 23, 128–137. 10.1016/j.ijdrr.2017.05.005

Baum, D. A., Bombacaceae, A., & Baum, D. A. (1995). A Systematic Revision of Adansonia (Bombacaceae) 82(3), 440–471. 10.2307/2399893

Baum, D. A., Small, R. L., & Wendel, J. F. (1998). Biogeography and flora evolution of Baobabs (Adansonia, Bombacaceae) as infered from multiple data sets. Systematic Biology, 47(2), 181–207. 10.1080/106351598260879

Beck, H. E., Wood, E. F., Pan, M., Fisher, C. K., Miralles, D. M., van Dijk, A. I. J. M., McVicar, T. R., and Adler, R. F. MSWEP V2 global 3-hourly 0.1° precipitation: methodology and quantitative assessment Bulletin of the American Meteorological Society 100(3), 473–500, 2019

Belmecheri, S., & Lavergne, A. (2020). Compiled records of atmospheric CO2 concentrations and stable carbon isotopes to reconstruct climate and derive plant ecophysiological indices from tree rings. Dendrochronologia, 63(August), 125748. 10.1016/j.dendro.2020.125748

Broccoli, A. J., Dahl, K. A., & Stouffer, R. J. (2006). Response of the ITCZ to Northern Hemisphere cooling. Geophysical Research Letters, 33(1), 1–4. 10.1029/2005GL024546

Burney, D. (1993). Late Holocene Environmental changes in arid southwestern Madagascar. Quaternary Research, 40, 98–106. 10.1006/qres.1993.1060

Burns, S. J., Godfrey, L. R., Faina, P., McGee, D., Hardt, B., Ranivoharimanana, L., & Randrianasy, J. (2016). Rapid human-induced landscape transformation in Madagascar at the end of the first millennium of the Common Era. Quaternary Science Reviews, 134, 92–99. 10.1016/j.quascirev.2016.01.007

Carrière, S.D., Chalikakis, K., Ollivier, C. et al. (2018). Sustainable groundwater resources exploration and management in a complex geological setting as part of a humanitarian project (Mahafaly Plateau, Madagascar). Environmental Earth Science 77, 734 10.1007/s12665-018-7909-1

Chase, B. M., & Meadows, M. E. (2007). Late Quaternary dynamics of southern Africa’s winter rainfall zone. Earth-Science Reviews, 84(3–4), 103–138. 10.1016/j.earscirev.2007.06.002

Chevalier, M., & Chase, B. M. (2015). Southeast African records reveal a coherent shift from high-to low-latitude forcing mechanisms along the east African margin across last glacial e interglacial transition. Quaternary Science Reviews, 125, 117–130. 10.1016/j.quascirev.2015.07.009

Chiang, J. C. H., & Bitz, C. M. (2005). Influence of high latitude ice cover on the marine Intertropical Convergence Zone. Climate Dynamics, 25(5), 477–496. 10.1007/s00382-005-0040-5

Christensen J.H., Hewitson B., Busuioc A., Chen A., Gao X., Held I., Jones R., Kolli R.K., Kwon W-T., Laprise R., Magana Rueda V., Mearns L., Menendez C.G., Raisanen J., Rinke A., Sarr A., Whetton P. (2007). Regional climate projections. In: Solomon S., Qin D., Manning M., Chen Z., Marquis M., Averyt A.B., Tignor M., Miller H.L. (eds) Climate change 2007: the physical science basis. Contribution of Working Group I to the Fourth Assessment Report of the Inter-governmental Panel on Climate Change. Cambridge University Press, Cambridge

Connolly-Boutin, L. & Smit, B. (2016). Climate change, food security, and livelihoods in sub-Saharan Africa. Regional Environmental Change, 16: 385–399. 10.1007/s10113-015-0761-x

Cook, E. R., Palmer, J. G., Cook, B. I., Hogg, A., & D’Arrigo, R. D. (2002). A multi-millennial palaeoclimatic resource from Lagarostrobos colensoi tree-rings at Oroko Swamp, New Zealand. Global and Planetary Change, 33(3–4), 209–220. 10.1016/S0921-8181(02)00078-4

Crossley, R., Davison-Hirschmann, S., Owen, R.B., Shaw, P.A. (1984). Lake level fluctuations during the last 2000 years in Malawi. J. Vogel (Ed.), Late Cainozoic Palaeoclimates of the Southern Hemisphere, A.A. Balkema, Rotterdam pp. 305–316. http://pascal-francis.inist.fr/vibad/index.php?action=getRecordDetail&idt=7268013

Donque, G. (1972). The climatology of Madagascar. In: Biogeography and ecology of Madagascar. R. Battistini and G. Richard-Vindard (eds). Junk, The Hague. 87–144. 10.1007/978-94-015-7159-3_3

Engelbrecht, F. A., Marean, C. W., Cowling, R. M., Engelbrecht, C. J., Neumann, F. H., Scott, L., Nkoana, R., O’Neal, D., Fisher, E., Shook, E., Franklin, J., Thatcher, M., McGregor, J. L., Van der Merwe, J., Dedekind, Z., & Difford, M. (2019). Downscaling Last Glacial Maximum climate over southern Africa. Quaternary Science Reviews, 226, 105879. 10.1016/j.quascirev.2019.105879

Faina, P., Burns, S. J., Godfrey, L.R., Crowley, B.E., Scroxton, N., McGee, D., Sutherland, M.R., & Ranivoharimanana, L. Comparing the paleoclimates of northwestern and southwestern Madagascar during the late Holocene: Implications for the role of climate in megafaunal extinction. Malagasy nature, 15 (). Retrieved from https://par.nsf.gov/biblio/10316730.

Farquhar, G., O’Leary, M. & Berry, J. (1982). On the Relationship between Carbon Isotope Discrimination and the Intercellular Carbon Dioxide Concentration in Leaves. Australian Journal of Plant Physiology. 9(2), 121–137. 10.1071/PP9820121

Ganzhorn, J. U. (1995). Cyclones over Madagascar: fate or fortune? Ambio. 24,124–125. http://www.jstor.org/stable/4314308

Gillett, N. P., Kell, T. D., & Jones, P. D. (2006). Regional climate impacts of the Southern Annular Mode. Geophysical Research Letters, 33(23), 1–4. 10.1029/2006GL027721

Grove, C. A., Zinke, J., Peeters, F., Park, W., Scheufen, T., Kasper, S., Randriamanantsoa, B., McCulloch, M. T., & Brummer, G. J. A. (2013). Madagascar corals reveal a multidecadal signature of rainfall and river runoff since 1708. Climate of the Past, 9(2), 641–656. 10.5194/cp-9-641-2013

Hahn, A., Schefuß, E., Groeneveld, J., Miller, C., & Zabel, M. (2021). Glacial to interglacial climate variability in the southeastern African subtropics (25–20° S). Climate of the Past, 17, 345–360. 10.5194/cp-2019-158

Hajdas, I., Hendriks, L., Fontana, A., & Monegato, G. (2017). Evaluation of Preparation Methods in Radiocarbon Dating of Old Wood. Radiocarbon, 59(3), 727–737. 10.1017/RDC.2016.98

Hall, G., Woodborne, S., & Pienaar, M. (2009). Rainfall control of the δ 13 C ratios of Mimusops caffra from KwaZulu-Natal, South Africa. The Holocene, 19(2), 251–260. 10.1177/0959683608100569

Hall, G., Woodborne, S., & Scholes, M. (2008). Stable carbon isotope ratios from archaeological charcoal as palaeoenvironmental indicators. 247, 384–400. 10.1016/j.chemgeo.2007.11.001

Hänke, H. & Barkmann, J. (2017). Insurance Function of Livestock: Farmer’s Coping Capacity with Regional Droughts in South-Western Madagascar. World Development. 96: 264–275. 10.1016/j.worlddev.2017.03.011

Hänke, H., Barkmann, J., Coral, C., Enforskaustky, E., & Marggraf, R. (2017). Social-ecological traps hinder rural development in Southwestern Madagascar. Ecology and Society, 22(1). 10.5751/ES-09130-220142

Haug, G., Hughen, K., Sigman, D., Peterson, L., & Ro, U. (2001). Southward Migration of the ITCZ holocene. Science, 293(August), 1304–1309. 10.1126/science.1059725

Helama, S., Timonen, M., Lindholm, M., Meriläinen, J., & Eronen, M. (2005). Extracting long-period climate fluctuations from tree-ring chronologies over timescales of centuries to millennia. International Journal of Climatology, 25(13), 1767–1779. 10.1002/joc.1215

Ho, C. H., Kim, J. H., Jeong, J. H., Kim, H. S., & Chen, D. L. (2006). Variation of tropical cyclone activity in the South Indian Ocean: El Nino-Southern Oscillation and Madden-Julian Oscillation effects. Journal of Geophysical Research-Atmospheres, 111(D22), D22101. Artn D22101\nDoi 10.1029/2006jd007289

Hoell, A., Funk, C., Zinke, J., & Harrison, L. (2017). Modulation of the Southern Africa precipitation response to the El Niño Southern Oscillation by the subtropical Indian Ocean Dipole. Climate Dynamics, 48(7–8), 2529–2540. 10.1007/s00382-016-3220-6

Hogg A.G., Hua Q., Blackwell P.G., Niu M., Buck C.E., Guilderson T.P., Heaton T.J., Palmer J.G. et al. (2013) SHCal13 Southern Hemisphere calibration, 0–50,000 cal yr BP. Radiocarbon. 55: 1889–1903. 10.2458/azu_js_rc.55.16783

Holmgren, K., Karlén, W., Lauritzen, S. E., Lee-Thorp, J. A., Partridge, T. C., Piketh, S., Repinski, P., Stevenson, C., Svanered, O., & Tyson, P. D. (1999). A 3000-year high-resolution stalagmite-based record of palaeoclimate for northeastern South Africa. Holocene, 9(3), 295–309. 10.1191/095968399672625464

Hoscilo, A., Balzter, H., Bartholomé, E., Boschetti, M., Brivio, P. A., Brink, A., Clerici, M., & Pekel, J. F. (2015). A conceptual model for assessing rainfall and vegetation trends in sub-Saharan Africa from satellite data. International Journal of Climatology, 35(12), 3582– 3592. 10.1002/joc.4231

Hua, Q., Barbetti, M., & Rakowski, A. (2013). Atmospheric Radiocarbon for the Period 1950– 2010. Radiocarbon, 55(4), 2059–2072. 10.2458/azu_js_rc.v55i2.16177

Huffman, T. N., & Woodborne, S. (2016). Archaeology, baobabs and drought: Cultural proxies and environmental data from the Mapungubwe landscape, southern Africa. Holocene, 26(3), 464–470. 10.1177/0959683615609753

Huffman, T.N. (2004) The archaeology of the Nguni past. Southern African Humanities. 16(1929): 79–111. https://hdl.handle.net/10520/EJC84742

Ingram, J. C., & Dawson, T. P. (2005). Inter-annual analysis of deforestation hotspots in Madagascar from high temporal resolution satellite observations. International Journal of Remote Sensing, 26(7), 1447–1461. 10.1080/01431160412331291189

IPCC (2021): Climate Change (2021) The Physical Science Basis. Contribution of Working Group I to the Sixth Assessment Report of the Intergovernmental Panel on Climate Change [Masson-Delmotte, V., Zhai, P., Pirani, A., Connors, S. L., Péan, C., Berger, S., Caud, N., Chen, Y., Goldfarb, L., Gomis, M. I., Huang, M., Leitzell, K., Lonnoy,., E., Matthews, J. A. R., Maycock, T. K., Waterfield, T., Yelekçi, O., Yu, R. & Zhou, B. (eds.)]. Cambridge University Press.

Johnson, T.C., Barry, S.L., Chan, Y., Wilkinson, P. 2001. Decadal record of climate variability spanning the past 700 yr in the Southern Tropics of East Africa. Geology 29 (1): 83–86. 10.1130/0091

Jury, M. R. (2016). Summer climate of Madagascar and monsoon pulsing of its vortex. Meteorology and Atmospheric Physics, 128(1), 117–129. 10.1007/s00703-015-0401-5

Jury, M. R., & Huang, B. (2004). The Rossby wave as a key mechanism of Indian Ocean climate variability. Deep Sea Research Part I: Oceanographic Research Papers, 51(12), 2123–2136. 10.1016/j.dsr.2004.06.005

Kaufmann, J. C. & Tsirahamba, S. (2006). Forests and Thorns: Conditions of Change Affecting Mahafale Pastoralists in Southwestern Madagascar. Conservation and Society. 4(2): 231– 261.

King, J., Anchukaitis, K.J., Allen, K. et al. Trends and variability in the Southern Annular Mode over the Common Era. Nat Commun 14, 2324 (2023). 10.1038/s41467-023-37643-1

Kuiper, M., Meijerink, G., & Eaton, D. (2007). Rural livelihoods: Interplay between farm activities, non-farm activities and the resource base. Science for Agriculture and Rural Development in Low-Income Countries, 77–95. 10.1007/978-1-4020-6617-7_5

Lechleitner, F. A., Breitenbach, S. F. M., Rehfeld, K., Ridley, H. E., Asmerom, Y., Prufer, K. M., Marwan, N., Goswami, B., Kennett, D. J., Aquino, V. V., Polyak, V., Haug, G. H., Eglinton, T. I., & Baldini, J. U. L. (2017). Tropical rainfall over the last two millennia: Evidence for a low-latitude hydrologic seesaw. Scientific Reports, 7(June 2016), 1–9. 10.1038/srep45809

Li, J., Xie, S.P., Cook, E. et al. (2011). Interdecadal modulation of El Niño amplitude during the past millennium. Nature Climate Change. 1, 114–118. 10.1038/nclimate1086

Liu, Z., Otto-Bliesner, Z., Kutzbach, J., Li, L. C. (2003). Shields Coupled climate simulation of the evolution of global monsoons in the Holocene. Journal of Climate, 16, 2472–2490. 10.1175/1520-0442(2003)016<2472:CCSOTE>2.0.CO;2

Loader, N. J., Robertson, I., Barker, A. C., Switsur, V. R., & Waterhouse, J. S. (1997). An improved technique for the batch processing of small wholewood samples to a-cellulose. Chemical Geology, 136, 313–317. 10.1016/S0009-2541(96)00133-7

MacDonald, G. M., & Case, R. A. (2005). Variations in the Pacific Decadal Oscillation over the past millennium. Geophysical Research Letters, 32(8), 1–4. 10.1029/2005GL022478

Macron, C., Pohl, B., Richard, Y., & Bessafi, M. (2014). How do tropical temperate troughs form and develop over Southern Africa? Journal of Climate, 27(4), 1633–1647. 10.1175/JCLI-D-13-00175.1

Mamalakis, A., Randerson, J.T., Yu, JY. et al. (2021). Zonally contrasting shifts of the tropical rain belt in response to climate change. Nature Climate Change 11, 143–151. 10.1038/s41558-020-00963-x

Manhique, A. J., Reason, C. J. C., Rydberg, L. & Fauchereau, N. (2011). ENSO and Indian sea surface temperatures with tropical temperate troughs over Mozambique and the southwest Indian Ocean. International Journal of Climatology. 31, 1–13. 10.1002/joc.2050

Mann, M. E., Zhang, Z., Rutherford, S., Bradley, R. S., Hughes, M. K., Shindell, D., Ammann, C., Faluvegi, G., & Ni, F. (2009). Global Signatures and Dynamical Origins of the Little Ice Age and Medieval Climate Anomaly. Science, 326(5957), 1256–1260. 10.1126/science.1177303

Mason, S. J. & Jury, M. R. (1997). ‘Climatic change and variability over Southern Africa: A Reflection on Underlying Processes’, Progress in Physical Geoggraphy. 21, 23–50. 10.1177/030913339702100103

Matsumoto, K. & Burney, D. A. (1994). Late Holocene environments at Lake Mitsinjo, northwestern Madagascar. Holocene, 4(1), 16–24. 10.1177/095968369400400103

McCarroll, D., & Loader, N. J. (2004). Stable isotopes in tree rings. Quaternary Science Reviews, 23(7–8), 771–801. 10.1016/j.quascirev.2003.06.017

Miller, C., Finch, J., Hill, T., Peterse, F., Humphries, M., Zabel, M., and Schefuß, E. (2019) Late Quaternary climate variability at Mfabeni peatland, eastern South Africa, ClimAte of the Past, 15,1153–1170, 10.5194/cp-15-1153-2019

Nakamura, N., Kayanne, H., Iijima, H., McClanahan, T.R., Behera, S.K., Yamagata, T., (2009). Mode shift in the Indian Ocean climate under global warming stress. Geophysical Research Letters. 36 10.1029/2009GL040590

Nash. (2017). Changes in Precipitation Over Southern Africa During Recent Centuries. Climate Science. 10.1093/acrefore/9780190228620.013.539

Nasri, M., & Modarres, R. (2009). Dry spell trend analysis of Isfahan Province, Iran. International Journal of Climatology, 29, 1430–1438. 10.1002/joc

Neukom, R., & Gergis, J. (2012). Southern Hemisphere high-resolution palaeoclimate records of the last 2000 years. The Holocene, 22(5), 501–524. 10.1177/0959683611427335

Nicholson, S. E., Klotter, D., & Dezfuli, A. K. (2012). Spatial reconstruction of semi-quantitative precipitation fields over Africa during the nineteenth century from documentary evidence and gauge data. Quaternary Research, 78(1), 13–23. 10.1016/j.yqres.2012.03.012

PAGES 2k Consortium. 2013. Continental-scale temperature variability during the past two millennia. Nature Geoscience. 6: 339–346. 10.1038/1849t

Patrut, A., Reden, K. F. Von, Danthu, P., Pock-tsy, J. L., Rakosy, L., Patrut, R. T., Lowy, D. A., & Margineanu, D. (2015). Nuclear Instruments and Methods in Physics Research B AMS radiocarbon dating of very large Grandidier’s baobabs (Adansonia grandidieri). Nuclear Inst. and Methods in Physics Research, B, 361, 591–598. 10.1016/j.nimb.2015.04.044

Onyeaka, H., Nwauzoma, U. M., Akinsemolu, A. A., Tamasiga, P., Duan, K., Al-Sharify, Z. T., & Siyanbola, K. F. (2024). The ripple effects of climate change on agricultural sustainability and food security in Africa. Food and Energy Security, 13, e567. 10.1002/fes3.567

Patrut, A., Woodborne, S., Von Reden, K. F., Hall, G., Patrut, R. T., Rakosy, L., Danthu, P., Pock-Tsy, J. M. L., Lowy, D. A., & Margineanu, D. (2017). The growth stop phenomenon of baobabs (Adansonia spp.) Identified by radiocarbon dating. Radiocarbon, 59(2), 435–448. 10.1017/RDC.2016.92

Pohlert, T. (2018). R Package “trends.” 18. https://cran.r-project.org/web/packages/trend/vignettes/trend.pdf

Putnam, A. E., & Broecker, W. S. (2017). Human-induced changes in the distribution of rainfall. Science Advances, 3(5), 1–14. 10.1126/sciadv.1600871

Railsback, L. B., Brook, G. A., Liang, F., Voarintsoa, N. R. G., Cheng, H., & Edwards, R. L. (2018). A multi-proxy climate record from a northwestern Botswana stalagmite suggesting wetness late in the Little Ice Age (1810–1820 CE) and drying thereafter in response to changing migration of the tropical rain belt or ITCZ. Palaeogeography, Palaeoclimatology, Palaeoecology, 506(April), 139–153. 10.1016/j.palaeo.2018.06.029

Ramanantsoa, J. D., Penven, P., Krug, M., Gula, J., & Rouault, M. (2018). Uncovering a new current: The Southwest Madagascar Coastal Current. Geophysical Research Letters, 45, 1930–1938. 10.1002/2017GL075900

Ratna, S. B., Behera, S., Ratnam, J. V., Takahasgi, K., & Yamagata, T. (2012) An index for tropical temperate troughs over southern Africa. Climate Dynamics, 41, 421–441. 10.1007/s00382-012-1540-8.

Razanamaro O., Rasoamanana E., Rakouth B., Randriamalala J.R., Rabakonadrianina E., Clément-Vidal A., Leong Pock Tsy J.M., Menut C., Danthu P. (2015). Chemical characterization of floral scents in the six endemic baobab species (Adansonia sp.) of Madagascar. Biochemical Systematics and Ecology, 60: 238–248 10.1016/j.bse.2015.04.005

Razanatsoa, E., Gillson, L., Virah-Sawmy, M., and Woodborne, S. (2021) Pollen records of the 14th and 20th centuries AD from Lake Tsizavatsy in southwest Madagascar. Palaeoecology of Africa, 35, pp. 309–315. 10.1201/9781003162766-20

Razanatsoa, E., Gillson, L., Virah-Sawmy, M., and Woodborne, S. (2022) Synergy between climate and human land-use maintained open vegetation in southwest Madagascar over the last millennium. The Holocene. 31(12). 10.1177/09596836211041731

Reason, C. J. C., & Mulenga, H. (1999). Relationships between South African rainfall and SST anomalies in the southwest Indian Ocean. International Journal of Climatology, 19(15), 1651–1673. 10.1002/(SICI)1097-0088(199912)19:15<1651::AID-JOC439>3.0.CO;2-U

Robertson, I., Loader, N., Froyd, C., Zambatis, N., Whyte, I., & S, W. (2006). The potential of the baobab (Adansonia digitata L.) as a proxy climate archive. Applied Geochemistry, 21, 1674–1680. 10.1016/j.apgeochem.2006.07.005

Russell, J. M. & Johnson, T. C. (2007). Little ice age drought in equatorial Africa: inter-tropical convergence zone migrations and El Niño southern oscillation variability. Geology. 35: 21–24. 10.1130/G23125A.1.

Russell, J. M., Verschuren, D., & Eggermont, H. (2007). Spatial complexity of “Little Ice Age” climate in East Africa: Sedimentary records from two crater lake basins in western Uganda. Holocene, 17(2), 183–193. 10.1177/0959683607075832

Sachs, J. P., Sachse, D., Smittenberg, R. H., Zhang, Z., Battisti, D. S., & Golubic, S. (2009). Southward movement of the Pacific intertropical convergence zone AD 1400-1850. Nature Geoscience, 2(7), 519–525. 10.1038/ngeo554

Saji, N. H., & Yamagata, T. (2003). Possible impacts of Indian Ocean Dipole mode events on global climate. Climate Research, 25(2), 151–169. 10.3354/cr025151

Schneider, T., Bischoff, T. & Haug, G. (2014). Migrations and dynamics of the intertropical convergence zone. Nature, 513, 45–53. 10.1038/nature13636

Scroxton, N., Burns, S. J., Mcgee, D., Hardt, B., Godfrey, L. R., Ranivoharimanana, L., & Faina, P. (2017). Hemispherically in-phase precipitation variability over the last 1700 years in a Madagascar speleothem record. 164. 10.1016/j.quascirev.2017.03.017

Serele, C., Pérez-Hoyos, A., & Kayitakire, F. (2020). Mapping of groundwater potential zones in the drought-prone areas of south Madagascar using geospatial techniques. Geoscience Frontiers, 11(4), 1403–1413. 10.1016/j.gsf.2019.11.012

Sigman, D. M., Hain, M. P., and Haug, G. H. (2010). The polar ocean and glacial cycles in atmospheric CO2 concentration, Nature, 466, 47–55, 10.1038/nature09149, 201

Stager, J. C., Ryves, D., Cumming, B. F., David Meeker, L., & Beer, J. (2005). Solar variability and the levels of Lake Victoria, East Africa, during the last millenium. Journal of Paleolimnology, 33(2), 243–251. 10.1007/s10933-004-4227-2

Svensson, A., Andersen, K. K., Bigler, M., Clausen, H. B., Dahl-Jensen, D., Davies, S. M., Johnsen, S. J., Muscheler, R., Rasmussen, S. O., Röthlisberger, R., Peder Steffensen, J., & Vinther, B. M. (2006). The Greenland Ice Core Chronology 2005, 15-42 ka. Part 2: comparison to other records. Quaternary Science Reviews, 25(23–24), 3258–3267. 10.1016/j.quascirev.2006.08.003

Tadross, M., Randriamarolaza, L., Rabefitia, Z., & Ki Yip, Z. (2008). Climate change in Madagascar; recent past and future. … DC (World Bank), February, 18. http://www.mediagrapher.org/gripweb/sites/default/files/disaster_risk_profiles/MadagascarClimateReport.pdf

Tambo, J.A. & Abdoulaye, T. (2013). Smallholder farmers’ perceptions of and adaptations to climate change in the Nigerian savanna. Regional Environmental Change 13, 375–388 10.1007/s10113-012-0351-0

Taylor, A. K., Berke, M. A., Castañeda, I. S., Koutsodendris, A., Campos, H., Hall, I. R., Hemming, S. R., LeVay, L. J., Sierra, A. C., & O’Connor, K. (2021). Plio-Pleistocene Continental Hydroclimate and Indian Ocean Sea Surface Temperatures at the Southeast African Margin. Paleoceanography and Paleoclimatology, 36(3), 1–18. 10.1029/2020pa004186

Thomas, D.S.G., Twyman, C., Osbahr, H. et al. (2007). Adaptation to climate change and variability: farmer responses to intra-seasonal precipitation trends in South Africa. Climatic Change 83, 301–322 10.1007/s10584-006-9205-4

Thompson, D. W. J., & Wallace, J. M. (2000). Annular modes in the extratropical circulation. Part I: Month-to-month variability. Journal of Climate, 13(5), 1000–1016. 10.1175/1520-0442(2000)013<1000:AMITEC>2.0.CO;2

Thompson, L. G., Mosley-Thompson, E., Davis, M. E., Lin, P. N., Henderson, K., & Mashiotta, T. A. (2003). Tropical glacier and ice core evidence of climate change on annual to millennial time scales. Climatic Change, 59(1–2), 137–155. 10.1023/A:1024472313775

Tierney, J. E., Smerdon, J. E., Anchukaitis, K. J. & Seager, R. (2013). Multidecadal variability in East African hydroclimate controlled by the Indian Ocean. Nature. 493: 389–392. 10.1038/nature11785

Tieszen, L. L. (1991). Natural variations in the carbon isotope values of plants: implications for archaeology, ecology and palaeoecology. Journal of Archaeological Science. 18: 227–248. 10.1016/0305-4403(91)90063-U

Trenberth, K.E. (1979). Interannual variability of the 500 mb zonal-mean flow in the Southern Hemisphere. Monthly Weather Reviews. 107, 1515–1524. 10.1175/1520-0493(1979)107<1515:IVOTMZ>2.0.CO;2

Tsen, E.W.J., Sitzia, T. and Webber, B.L. (2016). To core, or not to core: the impact of coring on tree health and a best-practice framework for collecting dendrochronological information from living trees. Biological Review, 91: 899–924. 10.1111/brv.12200

Tyson, P. D., Odada, E. O., & Partridge, T. C. (2001). Late quaternary environmental change in southern Africa. South African Journal of Science, 97(3–4), 139–150. https://hdl.handle.net/10520/EJC97298

Verschuren D. (2003). Lake-based climate reconstruction in Africa: progress and challenges. In: Martens K. (eds) Aquatic Biodiversity. Developments in Hydrobiology, vol 171. Springer, Dordrecht. 10.1007/978-94-007-1084-9_22

Verschuren, D., Laird, K. R., & Cumming, B. F. (2000). Rainfallanddroughtinequatorialeast Africa during the past 1, 100 years. Solar Cells, 403(January). 10.1038/35000179

Vigaud, N., Richard, Y., Rouault, M. & Fauchereau, N. (2007). Water vapour transport from the tropical Atlantic and summer rainfall in tropical southern Africa. Climate Dynamics, 28, 113–123. 10.1007/s00382-006-0186-9

Virah-Sawmy, M., Gillson, L., Gardner, C. J., Anderson, A., Clark, G., & Haberle, S. (2016). A landscape vulnerability framework for identifying integrated conservation and adaptation pathways to climate change: the case of Madagascar’s spiny forest. Landscape Ecology, 31(3), 637–654. 10.1007/s10980-015-0269-2

Voarintsoa, N. R. G., Wang, L., Railsback, L. B., Brook, G. A., Liang, F., Cheng, H., & Edwards, R. L. (2017). Multiple proxy analyses of a U/Th-dated stalagmite to reconstruct paleoenvironmental changes in northwestern Madagascar between 370 CE and 1300 CE. Palaeogeography, Palaeoclimatology, Palaeoecology, 469, 138–155. 10.1016/j.palaeo.2017.01.003

Von Heland, J., & Folke, C. (2014). A social contract with the ancestors-Culture and ecosystem services in southern Madagascar. Global Environmental Change, 24(1), 251–264. 10.1016/j.gloenvcha.2013.11.003

Wang, G., & Feng, X. (2012). Response of plants’ water use efficiency to increasing atmospheric CO2 concentration. Environmental Science & Technology, 46(16), 8610–8620. 10.1021/es301323m

Wang, L., Brook, G. A., Burney, D. A., Voarintsoa, N. R. G., Liang, F., Cheng, H., & Edwards, R. L. (2019). The African Humid Period, rapid climate change events, the timing of human colonization, and megafaunal extinctions in Madagascar during the Holocene: Evidence from a 2m Anjohibe Cave stalagmite. Quaternary Science Reviews, 210, 136–153. 10.1016/j.quascirev.2019.02.004

Watanabe, T. K., Watanabe, T., Yamazaki, A., Pfeiffer, M., & Claereboudt, M. R. (2019). Oman coral δ 18 O seawater record suggests that Western Indian Ocean upwelling uncouples from the Indian Ocean Dipole during the global-warming hiatus. Scientific Reports, 9(1), 1–9. 10.1038/s41598-018-38429-y

Weathering Risk (W.R). (2023). Climate Risk Profile for Southern Africa.

Wils THG, Robertson I, Woodborne S, Hall G, Koprowski M, Eshetu Z. 2016. Anthropogenic forcing increases the water-use efficiency of African trees. Journal of Quaternary Science 31, 386–390. 10.1002/jqs.2865

Woodborne S, Gandiwa P, Hall G, Patrut A, Finch J (2016) A Regional Stable Carbon Isotope Dendro-Climatology from the South African Summer Rainfall Area. PLoS ONE 11(7): e0159361. 10.1371/journal.pone.0159361

Woodborne, S., Hall, G., Robertson, I., Patrut, A., Rouault, M., Loader, N. J., & Hofmeyr, M. (2015). A 1000-year carbon isotope rainfall proxy record from South African baobab trees (Adansonia digitata L.). PLoS ONE, 10(5), 1–18. 10.1371/journal.pone.0124202

World Bank Group (2017). World Bank Annual Report (English). Washington, D.C. :. http://documents.worldbank.org/curated/en/143021506909711004/World-Bank-Annual-Report-2017

Yaro, J. A., Teye, J., & Bawakyillenuo, S. (2014). Local institutions and adaptive capacity to climate change/variability in the northern savannah of Ghana. Climate and Development, 7(3), 235–245. 10.1080/17565529.2014.951018

Zinke, J., Loveday, B. R., Reason, C. J. C., Kroon, D., Ocean, S. I., & Age, L. I. (2014). temperature variability in the Agulhas Current core region over the past 334. Scientific reports. pp. 1–8. 10.1038/srep04393

